# Population-scale immunoglobulin genomics links inherited variation to antibody repertoire organization

**DOI:** 10.64898/2026.06.05.730236

**Authors:** Ayelet Peres, Uddalok Jana, Oscar L. Rodriguez, Zachary M. Vanwinkle, Eric Engelbrecht, William S. Gibson, Kaitlyn Shields, Brandon Croslin, Steven Schultze, Chandrima Bharadwaj, Connor Murray, William D. Lees, Melissa L. Smith, Corey T. Watson, Gur Yaari

## Abstract

Human antibody repertoires emerge from inherited immunoglobulin variation interacting with V(D)J recombination and selection, yet how population-level germline diversity contributes to repertoire organization remains incompletely understood. We integrated long-read genomic data across the IGH, IGK, and IGL loci from 871 individuals, including 172 individuals with matched antibody repertoire sequencing. We identify extensive previously unrepresented genetic diversity and show that coding variation is embedded within a smaller set of recurrent RSS and leader sequence architectures. Integrating genomic and repertoire data further reveals genotype-associated differences in conditional segment usage that are not captured by conventional gene-usage analysis, and an association between IGK genotype similarity and exact CDR3 amino acid sharing across individuals. Together, these results reveal multiple levels at which inherited immunoglobulin variation is associated with the organization of expressed antibody repertoires. The underlying sequences and genomic and repertoire evidence are provided through the Human Unified Set of Alleles (HUSA), enabling genotype-aware analysis of immunoglobulin variation across diverse human populations.

**Significance statement:** Inherited variation in immunoglobulin genes influences antibody diversity, but the extent to which this variation shapes antibody repertoires remains poorly understood. By integrating population-scale genomic data with antibody repertoire sequencing, we reveal extensive previously uncharacterized genetic diversity, identify recurring relationships between coding and regulatory sequences, and demonstrate associations between inherited variation and repertoire organization. These findings provide a foundation for understanding how genetic diversity contributes to individual differences in antibody repertoires.

## Introduction

Antibody diversity is often described as a product of somatic processes. During B cell development, immunoglobulin (IG) heavy (IGH), kappa (IGK), and lambda (IGL) loci are rearranged through V(D)J recombination, and antigen-experienced B cells further diversify their receptors through somatic hypermutation (SHM) ^1,2^. These processes allow each person to generate an enormous and diverse repertoire of B-cell receptors and antibodies ^3,4^. Yet this diversity does not arise from a blank slate. It is built from inherited IG gene segments whose sequences, copy number, haplotypic organization, and local regulatory context vary across individuals ^5–9^. Therefore, the antibody repertoire observed after infection, vaccination, or immune challenge reflects both the germline architecture encoded in each individual’s genome and dynamic B cell processes ^10,11^. Understanding antibody repertoire variation therefore requires defining how germline IG variation is organized across individuals and populations and how this inherited variation contributes to the generation and selection of expressed antibody repertoires ^12^.

The biological importance of germline IG variation is increasingly clear. Coding and non-coding IG variants can influence the probability that specific V, D, and J genes are used, represented, or selected in expressed repertoires, thereby affecting the availability of antibody precursors, the sequence features available for antigen binding, and the paths available for affinity maturation ^6,7,11,13^. For example, IGHV1-2 germline variation has been linked to the frequency of VRC01-class HIV broadly neutralizing antibody precursors, a central consideration for germline-targeting vaccine design ^14–16^. Other IGH polymorphisms have been shown to influence anti-influenza antibody responses ^17^ and SARS-CoV-2 neutralizing antibodies ^18^. In these examples, allelic differences can alter residues or sequence contexts relevant to antibody-antigen interactions and can also shift the frequencies at which particular antibodies appear within circulating repertoires. Critically, functionally relevant alleles can vary substantially in frequency between human populations ^14,17,19^.

It is now widely understood that IG germline variation is not uniformly represented across populations or haplotypes, but the full extent of human diversity within the IG loci remains incompletely sampled. Repertoire-based haplotyping and long-read genomic analyses have revealed extensive allelic diversity and recurrent structural variation across IG loci ^8,9,19–21^. At the same time, probabilistic models of baseline repertoires are increasingly used to interpret antigen-driven responses and to predict repertoire features ^22–25^. The performance and interpretability of such models depend on accurate germline references and on a clearer understanding of the inherited features that structure repertoire formation. Thus, population-scale analyses of IG germline variation should consider not only coding alleles, but also the local sequence context in which each allele is inherited, so that coding variation can be interpreted together with linked recombination and regulatory features.

Identifying the germline features that shape recombination and repertoire composition therefore requires extending analysis beyond coding sequences to additional components of the IG loci that influence V(D)J recombination, such as recombination signal sequences (RSSs), other non-coding regulatory elements, and chromatin architecture ^26–29^. Variation in these features can influence whether a genomically encoded allele is observed in the expressed repertoire ^7,12^. Recently, gene-usage quantitative trait locus (guQTL) studies have tested whether inherited variants at these loci are statistically associated with variation in features of the expressed repertoire. These studies showed that single nucleotide and structural variants involving both coding and non-coding regions are associated with differences in V, D, and J gene usage, allele representation, CDR3 features, and broader genotype-dependent patterns of repertoire composition ^6,7^. Together, these observations argue that studies of expressed repertoires, vaccine responses, and immune-mediated phenotypes should account not only for the alleles carried by each individual, but also for the haplotype and population contexts in which those alleles occur. Defining IG variation at population scale is therefore critical to better understanding the relationships between the genome and the expressed repertoire. Each IG coding gene is inherited together with flanking and adjacent sequence features, including leader regions, recombination signal sequences (RSSs), and other linked regulatory elements. However, population-scale descriptions of variation in these gene-adjacent regions remain limited. Consistent with guQTL studies showing that variants across IG loci associate with gene and allele usage, CDR3 features, and broader repertoire composition ^6,7^, it is plausible that local sequence context and locus organization contribute to reproducible repertoire biases within individual haplotypes ^21^. Defining these relationships requires matched genomic and repertoire data that preserve physical linkage between coding sequences, adjacent non-coding features, and expressed receptor usage within individuals.

Here, we use population-scale IG long-read genomics and matched repertoire sequencing to test whether inherited coding variation is organized with linked regulatory sequence features and whether this organization is reflected in expressed antibody repertoires. We analyzed long-read genomic data for the IGH, IGK, and IGL loci from the Human Pangenome Reference Consortium (HPRC) ^30,31^ together with targeted IG long-read genomic data from a large collection of samples of diverse genetic ancestries, including a subset of donors with matched genomic-repertoire datasets. This design allowed us to distinguish allele presence in the genome from allele recovery in expressed repertoires, while preserving linkage between coding sequences, RSSs, leader regions, and repertoire usage within individuals. Using this framework, we show that human IG allelic variation is extensive across the human population, exhibiting population- and locus-specific genetic signatures. Additionally, we observe that IG alleles are best viewed not solely as isolated coding variants or gene-level units, but instead as linked coding regions, RSSs, and leader regions that represent inherited sequence blocks. We characterize these local sequence architectures and separately examine how inherited IG variation is associated with conditional V(D)J segment-pair structure and exact CDR3 amino acid sharing. Together, these analyses support a genomic-context-aware view of antibody repertoire formation in which inherited coding, recombination-signal, and leader variation provide a linked genomic context for repertoire structure. The Human Unified Set of Alleles (HUSA) provides the evidence-linked dataset underlying these analyses and a foundation for future genotype-aware studies of immune receptor variation. HUSA is available through VDJbase ^32,33^ and the AKC portal ^34^, with allele sequences linked to supporting evidence and associated metadata. VDJbase also provides interactive dashboards for exploring alleles and genes represented in HUSA and their associated guQTL results. Germline sets incorporating HUSA sequences will be published on OGRDB ^33^.

## Results

We first generated an integrated genomic and repertoire dataset to determine how inherited IG variation is organized across coding and linked regulatory sequence features at population scale. The study combined three complementary data sources spanning the IGH, IGK, and IGL loci: targeted long-read genomic data, publicly available HPRC long-read whole-genome data, and a matched cohort with targeted long-read genomic sequencing and AIRR-seq from the same individuals. This design allowed us to separate inherited allele presence from repertoire-based allele recovery while preserving allele-level links to RSS and leader sequence context.

### Population-scale immunoglobulin genomics resolves inherited variation across heavy and light chain loci

We constructed an evidence-linked allele dataset by integrating genomic and repertoire data across the IGH, IGK, and IGL loci. The study combines data from three sources, DS1, DS2, and DS3 (Table 1; Figure 1A). DS3 comprised matched targeted IGH/K/L PacBio high-fidelity (HiFi) long-read genomic sequencing and short-read 5’RACE IgM/IgK/IgL AIRR-seq data from the same individuals in a mixed-ancestry cohort (*n* = 172). DS1 comprised newly generated targeted HiFi long-read sequencing data from 541 individuals collected as part of the 1000 Genome Project (1KGP) ^35^, representing five super-populations: African (AFR), Admixed American (AMR), East Asian (EAS), European (EUR), and South Asian (SAS). DS2 comprised publicly available long-read whole-genome sequencing (WGS) data from 158 individuals generated by the Human Pangenome Reference Consortium (HPRC), also representing AFR, AMR, EAS, EUR, and SAS populations. The breakdown of cohorts by self-reported ancestry (DS3) and super-population (DS1 and DS2) is summarized in Table 1.

**Table 1:** Datasets analyzed in this study.

| **Dataset label** | **Cohort source** | **Data type** | **Retained n (IGH/IGK/IGL)** | **Ancestry/population groups** |
|---|---|---|---|---|
| DS1 | 1KGP samples | Targeted long-read genomic | 496/496/496 | AFR, AMR, EAS, EUR, SAS |
| DS2 | HPRC samples | Long-read whole-genome | 152/144/158 | AFR, AMR, EAS, EUR, SAS |
| DS3 | R24 samples | Targeted long-read genomic + AIRR-seq | 172/172/172 | AFR, AMR, EAS, EUR, SAS, Mixed |

**Figure 1:**
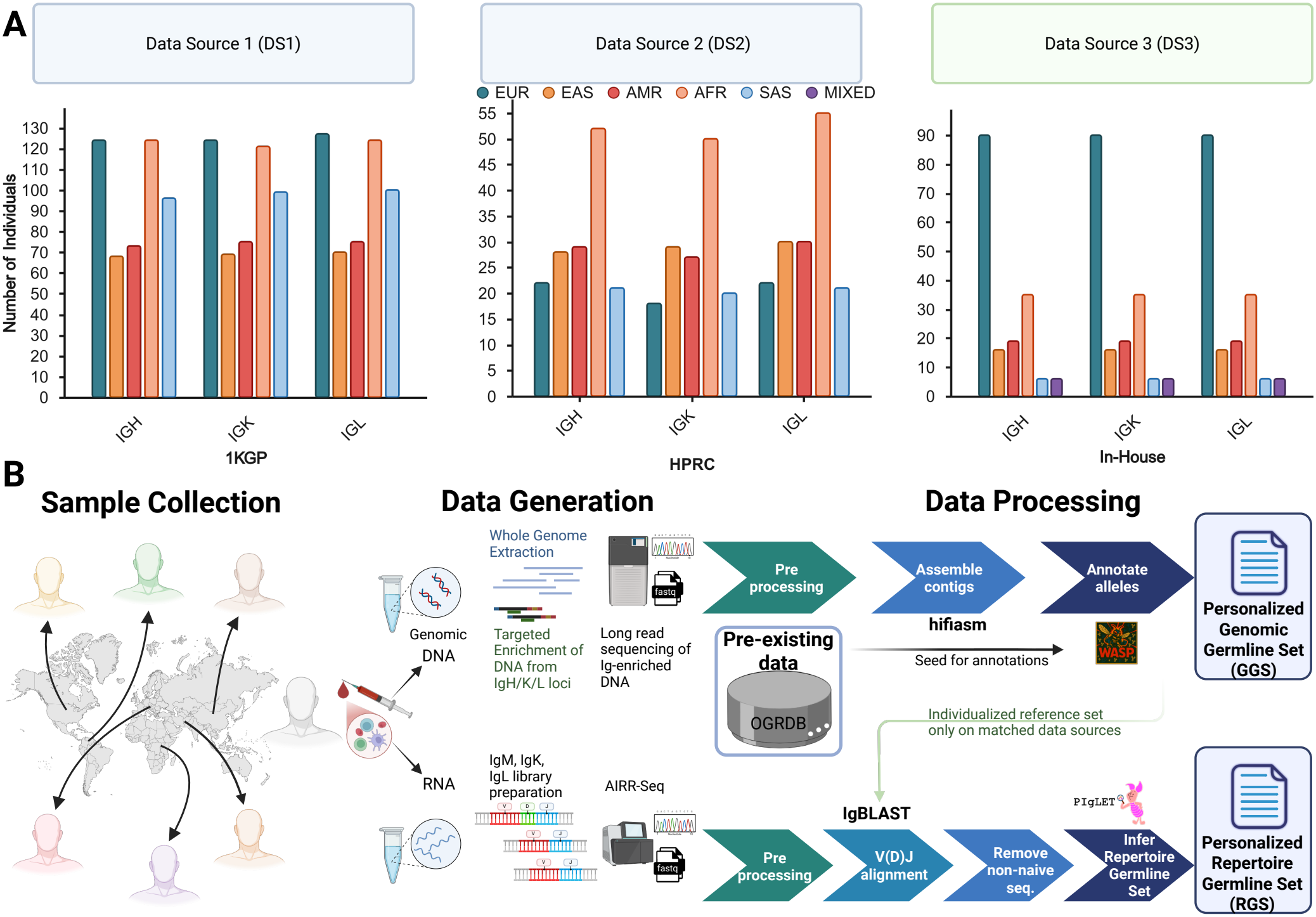
Data sources and processing workflow used to construct genomic and repertoire germline sets. (A) Number of individuals retained for each locus (IGH, IGK, and IGL), stratified by ancestry group, across DS1, DS2, and DS3 (Table 1). (B) Overview of sample collection, data generation, and processing. Genomic DNA from the IGH, IGK, and IGL loci was target-enriched and sequenced using long-read sequencing, while matched IgM, IgK, and IgL AIRR-seq libraries were generated from RNA. Genomic reads were preprocessed, assembled with hifiasm, and annotated with WASP using OGRDB as the seed reference to generate individualized GGSs. For DS3 samples, individualized genomic references were used for IgBLAST-based V(D)J alignment, filtering of non-naive sequences, and inference of personalized RGSs with PIgLET ^38^. DS1 and DS2 contributed genomic evidence only. Figure was created with https://BioRender.com.

All samples from DS1 and DS3 were required to have mean HiFi read coverage of at least 20*×* across the IGH, IGK, and IGL loci. These targeted HiFi reads were used to generate *de novo* diploid assemblies for the IGH, IGK, and IGL loci using Hifiasm ^36^, followed by gene annotation with the reference-guided WASP workflow ^37^. For DS2, we extracted assembly contigs spanning the IGH, IGK, and IGL loci and applied the same reference-guided annotation strategy.

For DS3, individualized genomic gene and allele annotations were used to derive genomic germline sets (GGSs), which then served as personalized references for repertoire alignment and repertoire germline set (RGS) inference. Additional subject-level filters were applied to support matched genomic-repertoire analyses (see Methods). This design preserved the distinction between inherited alleles detected in the genome and alleles recovered from expressed repertoire data in the HUSA analyses described below.

### Human immunoglobulin diversity shows population and locus specific structure

We next integrated genomic allele calls from DS1, DS2, and DS3 with the baseline reference to construct the HUSA. The baseline reference comprised all alleles from OGRDB together with 28 additional alleles from selected IG genes that were not included in the original OGRDB set publication. In total, HUSA contains 1,447 alleles across V, D, and J gene segments, including 1046 alleles absent from the baseline reference (Figure 2A). All alleles entering HUSA were required to satisfy conservative support criteria consistent with policies of the International Union of Immunological Societies TR-IG Nomenclature Review Committee (see Methods).

**Figure 2:**
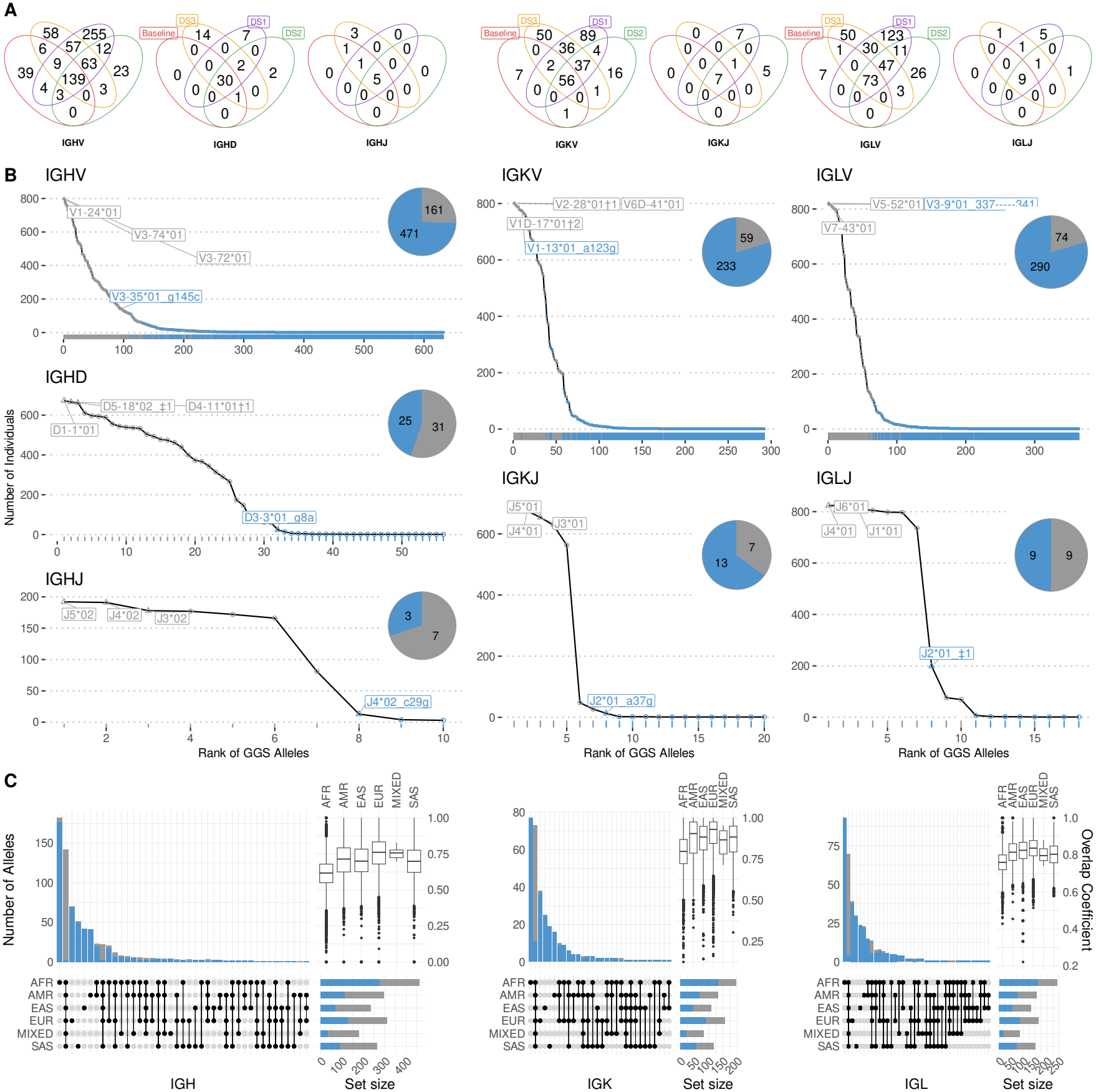
HUSA expands the population-scale IG allele set and reveals ancestry- and locus-specific sharing patterns. (A) Venn diagrams showing overlap among the baseline reference, DS3, DS1, and DS2 allele sets, stratified by gene segment. (B) Rank distributions of HUSA alleles across individuals in the genomic datasets. Only alleles detected in at least one individual were included. The x-axis indicates allele rank, and the y-axis shows the number of individuals carrying each allele. Each point represents an allele, with color indicating baseline or non-baseline status. The most widely shared alleles are annotated, and pie charts show the proportion of baseline and non-baseline alleles in each germline set. Daggers (*†*) mark labels that collapse several co-listed alleles into the first allele shown: in IGHD, *†*1 = IGHD4-11*01 and IGHD4-4*01/IGHD4-11*01, and *†*2 = IGHD5-18*02 a4g t8c g10-t11-g12-c17t a19g and IGHD5-5*01; in IGKV, *†*1 = IGKV1-39*01, IGKV1D-39*01 and IGKV1-39*01/IGKV1D-39*01, and *†*2 = IGKV2-28*01, IGKV2D-28*01 and IGKV2-28*01/IGKV2D-28*01. (C) UpSet plots showing the overlap of genomic alleles across ancestry groups, stratified by locus. Horizontal bars indicate the total number of alleles identified within each ancestry group, and vertical bars represent the number of alleles shared across ancestry combinations. Bars are colored by baseline or non-baseline status. Insets show within-ancestry overlap coefficients of GGS alleles for the IGH, IGK, and IGL loci.

Non-baseline alleles accounted for 72.3% of HUSA, corresponding to a 260.8% increase relative to the 401 alleles in the baseline reference. Expansion was observed across all loci and segment classes, with the largest absolute increases among V genes. IGHV contained 671 alleles, including 471 non-baseline alleles (70.2%); IGKV contained 300 alleles, including 234 non-baseline alleles (78.0%); and IGLV contained 372 alleles, including 291 non-baseline alleles (78.2%). Non-baseline alleles were also identified among D and J genes, including 25 of 56 IGHD alleles (44.6%), 3 of 10 IGHJ alleles (30.0%), 13 of 20 IGKJ alleles (65.0%), and 9 of 18 IGLJ alleles (50%).

We compared HUSA with both the baseline reference and IMGT (Sup. Figure 1). HUSA expanded beyond both resources across all V, D, and J segments while retaining substantial overlap with established reference alleles. We additionally summarized HUSA diversity using allele similarity clusters (ASCs), which group closely related coding-region sequences ^38^. Allele content varied substantially among ASCs across V, D, and J segment classes, with clusters containing different combinations of genomic and repertoire-supported alleles (Sup. Figure 2). We then examined allele prevalence across individuals in the genomic datasets (Figure 2B). Baseline alleles were generally observed in the greatest number of individuals, whereas most non-baseline alleles were observed in fewer individuals. However, non-baseline alleles were not limited to singletons: 454 were present in at least 2 individuals, 164 in at least 10 individuals, 49 in at least 40 individuals, and 20 in at least 90 individuals.

We next summarized the sharing of alleles between ancestry groups using UpSet plots colored by baseline versus non-baseline allele status, together with overlap coefficients within-ancestry (Figure 2C). African ancestry samples contained the largest number of observed alleles across all three loci, with 463 alleles in IGH, 198 in IGK, and 243 in IGL. In all populations, IGH harbored the greatest number of observed alleles. Although unique alleles were identified in all populations, the largest set of population-specific alleles was observed in AFR samples. Because raw allele counts scale with sample size, we based the diversity comparison on the per-pair within-ancestry overlap coefficient, which is independent of group size. This coefficient was calculated as the fraction of the smaller allele set shared by the other individual. By this measure, African ancestry samples showed the lowest median overlap across all three loci, with median coefficients of 0.617 for IGH, 0.796 for IGK, and 0.76 for IGL, compared with 0.761, 0.930, and 0.837 in European ancestry samples. Additionally, IGH had consistently lower median coefficients than IGK across all populations.

Together, these results showed that the inherited sequence space available for antibody repertoire formation is substantially broader than represented by current baseline references. Importantly, many non-baseline alleles were observed across multiple individuals rather than as isolated singletons, indicating that unresolved IG variation includes population-level alleles. This expanded view of genomic diversity allowed us to move beyond allele discovery and ask which alleles are recovered in expressed repertoires and how their coding sequences are organized with linked RSS and leader sequence contexts.

### Coding variation recovered in expressed repertoires occurs within diverse genomic contexts

After defining the expanded HUSA allele set, we examined the subset recovered from expressed repertoires in DS3. From this point onward, DS3 analyses were restricted to samples that passed the required repertoire-depth threshold (see Methods). Here, “expressed” refers to alleles inferred from productive AIRR-seq data using subject-specific genomic references and PIgLET-optimized inclusion thresholds, rather than to direct measurement of transcription from unrearranged germline loci (See Method section AIRR-seq Processing, Novel Allele Inference, and RGS Inference) ^38^. RGS inference identified 658 expressed HUSA alleles, nearly half of which were absent from the baseline reference (326 of 658, 50%; Figure 3A). The non-baseline fraction was comparable between light chains and IGH, accounting for 60.6% of IGK alleles and 54.4% of IGL alleles, compared with 41.3% of IGH alleles. These findings show that many non-baseline HUSA alleles are present not only in genomic data but are also recovered in productive repertoires, highlighting the importance of expanded reference sets for expressed repertoire analysis.

**Figure 3:**
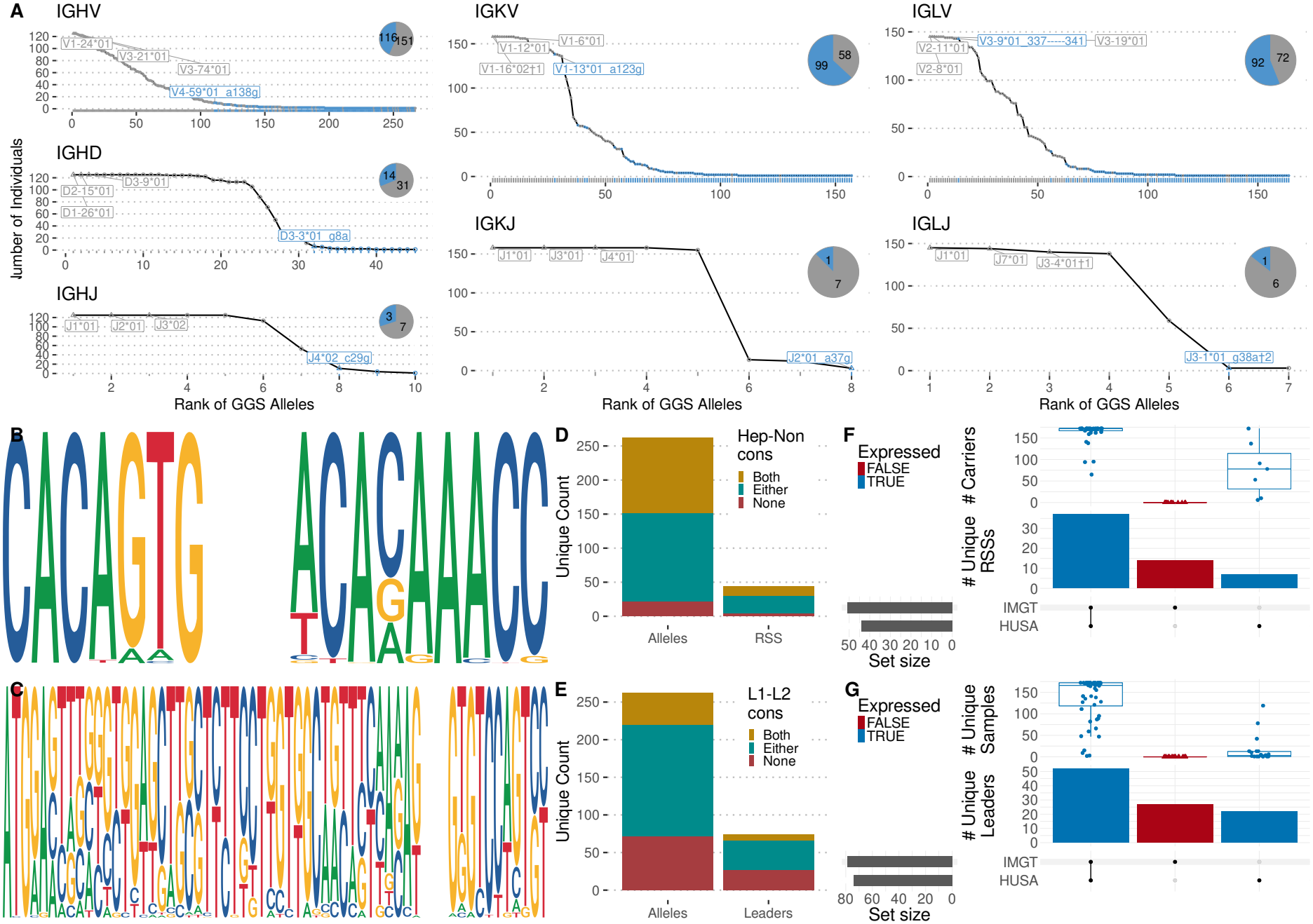
Expressed HUSA alleles are linked to RSS and leader sequence contexts. (A) Rank distributions of expressed HUSA alleles inferred from repertoire genotype sets in DS3. Each point represents an allele detected in at least one individual, with color indicating baseline or non-baseline status. The most widely shared alleles are annotated, and pie charts show the proportion of baseline and non-baseline alleles within each expressed germline set. Daggers (*†*) mark labels that collapse several co-listed alleles into the first allele shown: in IGLJ, *†*1 = IGLJ3-401, IGLJ3-201, IGLJ3-101, IGLJ3-301 g2t t3g a7g, IGLJ201 g2t t3g a7g, and IGLJ302/IGLJ3-101/IGLJ3-401/IGLJ3-201; *†*2 = IGLJ201, IGLJ3-301, IGLJ3-101 t2g g3t g7a, IGLJ3-401 t2g g3t g7a, and IGLJ201/IGLJ3-301; and *†*3 = IGLJ3-101 g38a and IGLJ3-401 g38a. (B-C) Sequence logos for IGHV RSS heptamer and nonamer motifs (B) and the combined IGHV leader sequence (C), based on unique allele-associated sequences from the DS3 genomic export used for matched repertoire analyses. (D-E) Counts of unique expressed IGHV alleles and associated RSS variants stratified by heptamer-nonamer consensus status (D), and counts of unique IGHV alleles and leader sequences stratified by leader 1-leader 2 consensus status (E). Consensus categories indicate whether both, either, or neither motif or leader segment matched the corresponding IGHV consensus. (F-G) UpSet plots comparing IGHV RSS variants (F) and IGHV leader sequences (G) represented in HUSA and IMGT. Bars show source-specific or shared sequence counts; boxplots show the number of DS3 carriers for each sequence; color indicates whether the associated allele was observed in expressed repertoire data.

We next examined the non-coding sequence contexts linked to expressed alleles in the same matched samples. For IGHV, sequence logos showed a strong heptamer consensus within RSSs, greater variability in the nonamer, and broad diversity across leader sequences (Figure 3B-C). Among expressed IGHV alleles, RSS variants were classified according to whether both, either, or neither the heptamer and nonamer matched the IGHV consensus. Leader sequences were classified analogously based on whether leader exon 1 and exon 2 matched the corresponding consensus sequences (Figure 3D-E).

Comparison with IMGT showed that IGHV RSS variants and leader sequences included both shared and HUSA-specific sequence classes. Importantly, a subset of HUSA-specific sequence classes were linked to alleles detected in expressed repertoire data, revealing additional linked sequence contexts that were not previously associated with alleles recovered from productive repertoires (Figure 3F-G). Comparable analyses across additional loci showed that regulatory sequence diversity was not restricted to IGHV, but extended across V, D, and J gene classes (Sup. Figure 3; Sup. Figure 4).

**Figure 4:**
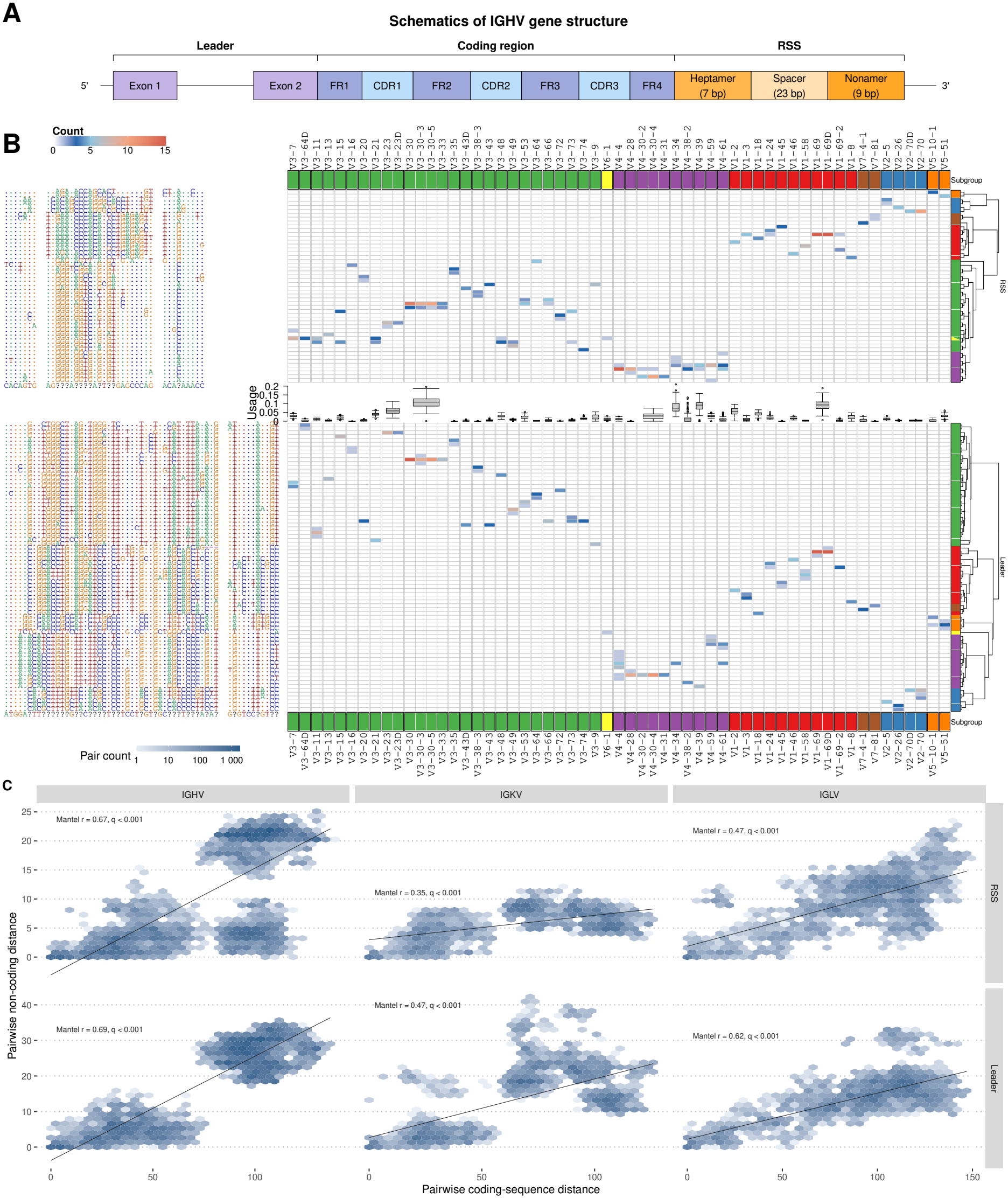
IGHV coding-region, RSS, leader, and repertoire-usage variation. (A) Schematic of an IGHV gene, including exon 1, exon 2, coding-region features, and the downstream recombination signal sequence (RSS) composed of the heptamer, 23-bp spacer, and nonamer. (B) Heatmaps connecting IGHV coding-region ASCs with RSS sequences, relative usage in expressed repertoires, and leader exon 1 and exon 2 sequence variants. Cell color denotes the number of alleles with each variant combination. Rows are hierarchically clustered, and columns are ordered by chromosomal position within subgroup. (C) Relationship between pairwise coding-region distance and pairwise RSS, exon 1, and exon 2 distance within IGHV, IGKV, and IGLV. Each panel shows a two-dimensional density with a linear trend. Mantel correlation coefficients and Benjamini-Hochberg-corrected *q* values are shown.

Distinguishing HUSA from a coding-allele centric germline catalog, these results expand knowledge of regulatory element diversity associated with expressed IG alleles and enable analysis of the local genomic contexts in which repertoire-associated alleles are inherited.

### V-gene coding diversity is linked to recurrent RSS and leader sequence contexts

Having identified substantial coding, RSS, and leader sequence diversity across the IG loci, we next examined how these sequence features are organized relative to one another. We first focused on V genes, where coding regions could be linked directly to their associated RSS and leader sequences. Figure 4A illustrates this organization for IGHV, with the leader upstream of the coding region and the 23-bp RSS downstream. Mapping these features across IGHV showed that coding diversity substantially exceeded the diversity of the linked sequence context (Figure 4B). Among 262 IGHV coding alleles with linked genomic annotations, we identified only 44 unique complete RSSs and 74 unique leader sequences. Similar compression was observed in IGKV, where 155 coding alleles were associated with 34 RSSs and 60 leaders, and in IGLV, where 163 coding alleles were associated with 43 RSSs and 72 leaders (Sup. Figure 5). Thus, expansion of V-gene coding diversity does not correspond to a one-to-one expansion of linked RSS and leader diversity.

**Figure 5:**
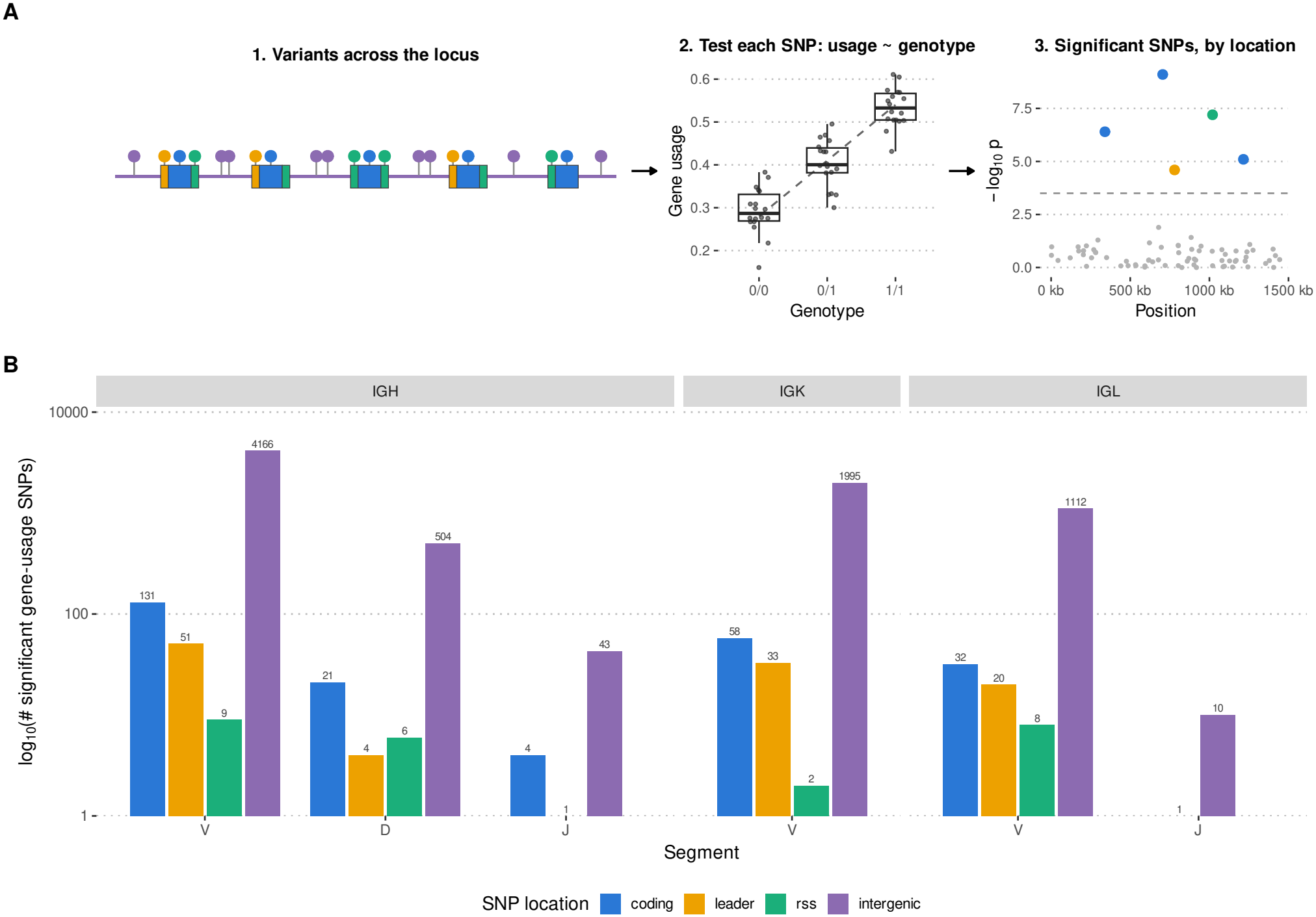
Genetic variants across immunoglobulin loci are associated with segment usage. (A) Schematic representation of the gene-usage QTL (guQTL) analysis. Variants across each IG locus were tested for association with segment usage as a function of genotype, and significant associations were mapped to their positions within the corresponding locus. (B) Summary of significant guQTL SNPs across IGH, IGK, and IGL, stratified by V, D, and J segment and SNP location. Bars show the number of significant gene-usage SNPs on a log_10_ scale, with raw counts indicated above each bar. SNPs are classified according to their location within coding regions, leader sequences, recombination signal sequences (RSSs), or intergenic regions.

The ASC-level maps further showed extensive reuse of flanking sequence contexts across coding groups. In IGHV, 13 RSS variants were associated with more than one gene, with a single RSS shared across seven genes. Although most sharing occurred among related genes, it was not restricted to individual V subgroups. IGHV6-1, for example, carried an RSS identical to that observed for IGHV3-7, IGHV3-11, IGHV3-21, IGHV3-48, and IGHV3-64D, despite coding-region distances of 104–117 nucleotides between IGHV6-1 and these IGHV3 genes. Leader sequences were also reused across coding backgrounds, with one IGHV leader shared by five genes. These patterns indicate that V-gene coding alleles are distributed across a smaller recurrent set of linked RSS and leader contexts rather than forming unique coding-RSS-leader combinations.

Relative repertoire usage also varied across these coding groups, including among ASCs sharing recurrent RSS or leader sequence contexts (Figure 4B; Sup. Figure 5), indicating that coding and flanking sequence relationships occur across a broad range of repertoire abundances.

RSS organization extended beyond V genes. IGHD showed distinct sequence structure at both flanking RSSs, with the 5′ and 3′ RSSs forming different patterns across D segments (Sup. Figure 6A). The 5′ IGHD heptamer was approximately twice as variable per site as the 3′ heptamer (0.218 versus 0.104), with 15 unique 5′ heptamers compared with 8 at the 3′ RSS, whereas greater spacer and nonamer diversity was observed at the 3′ side. RSS variation was also evident across IGHJ, IGKJ, and IGLJ segments (Sup. Figure 6B–D), demonstrating that recurrent and segment-specific RSS sequence structure is not restricted to V genes.

**Figure 6:**
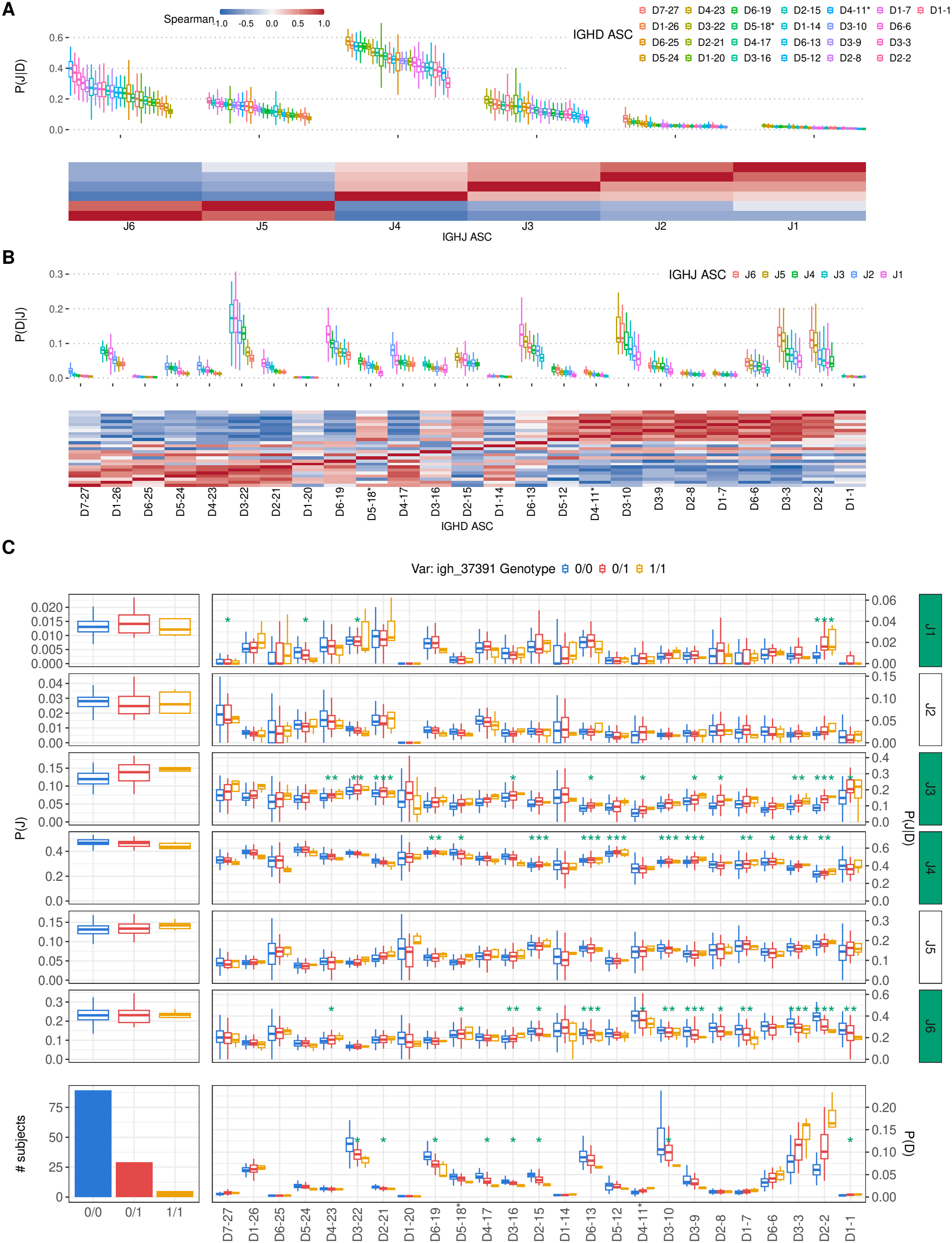
Conditional IGH segment usage and genotype-associated variation in D–J pairing. (A) *P* (*J* | *D*), showing IGHJ ASC usage conditional on IGHD ASC. Boxplots show individual-level conditional usage distributions, and the heatmap shows Spearman correlations between IGHJ ASC usage profiles. ( B) *P* ( *D* | *J*), showing IGHD ASC usage conditional on IGHJ ASC. Boxplots show individual-level conditional usage distributions, and the heatmap shows Spearman correlations between IGHD ASC usage profiles. Alleles were grouped into allele

Despite the reuse of individual regulatory sequences, coding and linked sequence variation remained strongly structured among V genes. Pairwise coding-region distance was significantly correlated with both RSS and leader distance across all three V loci (all *q <* 0.001; Figure 4C). In IGHV, coding distance correlated with RSS distance at *r* = 0.67 and leader distance at *r* = 0.69. Corresponding correlations were *r* = 0.35 and 0.47 in IGKV, and *r* = 0.47 and 0.62 in IGLV. Importantly, coding–RSS concordance persisted when comparisons were restricted to genes within the same V subgroup. Significant within-subgroup correlations ranged from *r* = 0.29 to 0.74 in IGHV, *r* = 0.32 to 0.88 in IGKV, and *r* = 0.36 to 0.91 in IGLV among sufficiently represented subgroups. The association between coding and RSS sequence therefore reflects structure among closely related genes and is not explained solely by sequence differences between V-gene subgroups.

We next decomposed RSS variation into the conserved heptamer and nonamer motifs and the intervening spacer across the complete set of 12- and 23-bp RSS classes (Sup. Figure 7). The distance heatmaps showed markedly different organization of these components across V, D, and J segments. Heptamer-plus-nonamer sequences were broadly conserved across many segment groups, whereas spacer sequences showed substantially greater divergence and stronger segment- and subgroup-specific structure. Quantitatively, complete RSS distance was more strongly correlated with spacer distance than with heptamer-plus-nonamer distance. For IGHV, IGKV, and IGLV, complete-RSS versus spacer correlations were *r* = 0.95, 0.88, and 0.95, respectively, compared with *r* = 0.73, 0.46, and 0.50 for the heptamer-plus-nonamer motifs. Per-site diversity was similarly lowest in the heptamer, ranging from 0.06 to 0.11 across the V loci, and substantially higher in the spacer, ranging from 0.33 to 0.47. These results show that much of the sequence divergence distinguishing complete RSS contexts is carried by the spacer, while the recombination-critical motifs remain comparatively constrained.

**Figure 7:**
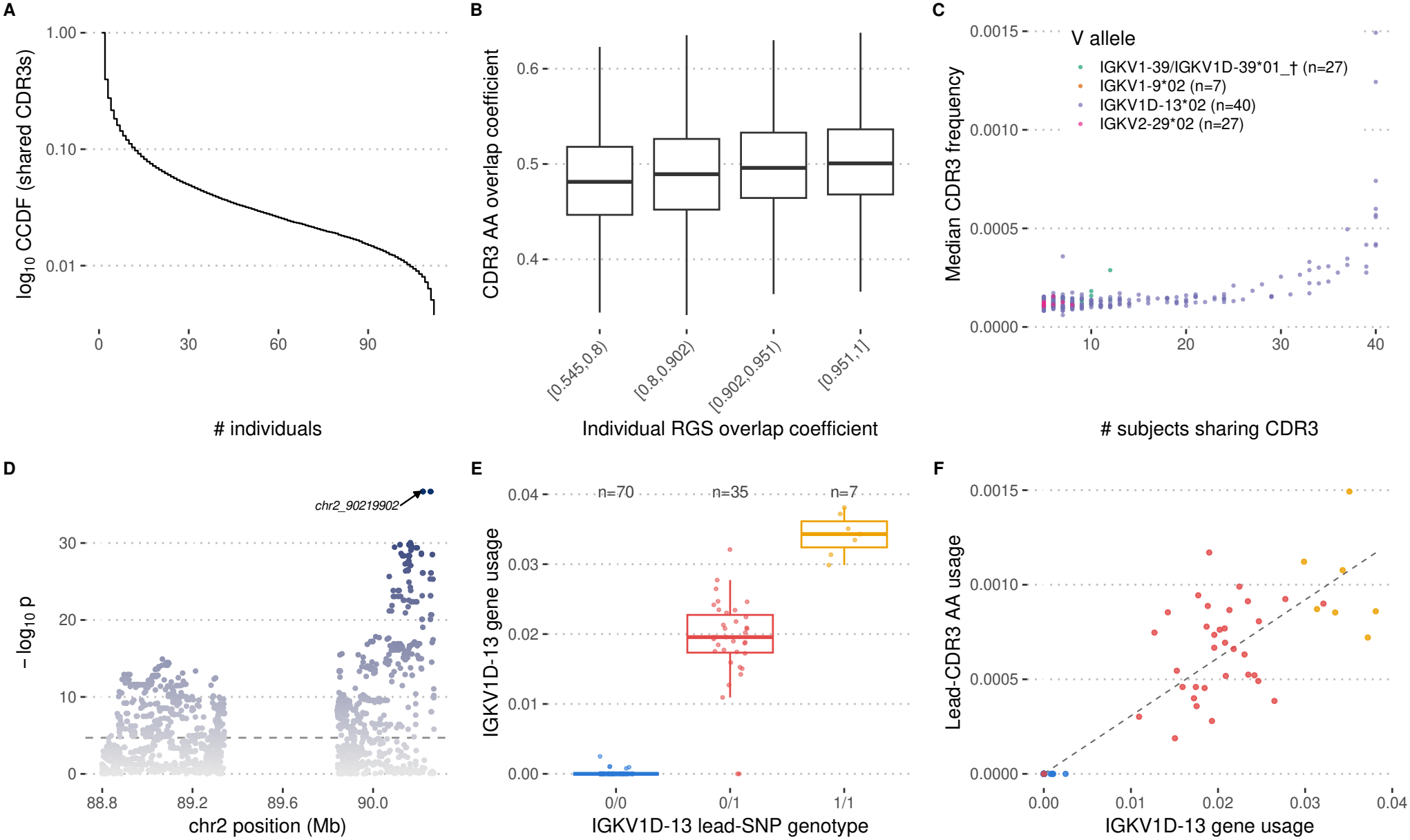
IGK CDR3 sharing is associated with repertoire genotype and inherited variation in IGKV1D-13 usage. (A) Complementary cumulative distribution function (CCDF) of exact IGK CDR3 amino acid (AA) sequences shared across individuals. The x-axis shows the number of individuals sharing a CDR3 AA sequence, and the y-axis shows the log_10_ cumulative probability of sequences shared by at least that number of individuals. (B) Relationship between repertoire genotype set (RGS) overlap and IGK CDR3 AA overlap across pairs of individuals. Individual pairs were grouped by RGS overlap coefficient, and boxplots show the corresponding distributions of exact CDR3 AA overlap. (C) IGK V allele-specific shared CDR3 AA sequences. Each point represents an exact CDR3 AA sequence observed in at least five individuals and assigned to a single IGK V allele. The x-axis shows the number of subjects sharing the CDR3 AA sequence, and the y-axis shows its median within-individual frequency. Only V alleles observed in no more than 60% of individuals were included. The legend label IGKV1-39/IGKV1D-39*01 *†* denotes the co-listed alleles IGKV1-3901 c29t c306t 315gtgtg319 c330a and IGKV1D-3901 c29t c306t 315gtgtg319 c330a. (D) Manhattan plot of SNP associations with IGKV1D-13 gene usage across the IGK locus. The dashed line indicates the significance threshold, and the lead variant, *chr2 90219902*, is labeled. (E) IGKV1D-13 gene usage stratified by lead-SNP genotype. Boxplots show usage among individuals with geno-types 0/0, 0/1, and 1/1, with the number of individuals in each genotype group indicated above the corresponding boxplot. (F) Relationship between IGKV1D-13 gene usage and the within-individual frequency of the most widely shared IGKV1D-13-associated CDR3 AA sequence identified in (C). Points are colored by lead-SNP genotype, and the dashed line shows the fitted relationship between gene usage and CDR3 AA usage.

Together, these analyses reveal a layered organization of IG germline sequence variation. Extensive V-gene coding diversity is embedded within a smaller recurrent set of RSS and leader contexts, while RSS sequence structure extends across V, D, and J segments and differs between the two flanks of IGHD genes. At the same time, coding and linked RSS variation retain significant concordance even among closely related V genes, and variation in the complete RSS is driven disproportionately by the spacer rather than the conserved heptamer and nonamer motifs.

### Genetic variation is associated with single and conditional IG segment usage

Genetic variation within the IG loci has previously been associated with inter-individual differences in gene usage ^6,7^. Following the same guQTL framework, we characterized associations between IG variant genotypes and individual gene segment usage across IGH, IGK, and IGL (Figure 5A). The IGK and IGL guQTLs shown here were previously reported using the same dataset ^7^ and are included alongside IGH to compare the genomic distribution of usage-associated variants across the three loci within the dataset used here.

Significant guQTLs were discovered in both coding and non-coding regions of the IG loci (Figure 5B). For V segments, IGH included 131 significant coding, 51 leader, and 9 RSS variants; IGK included 58, 33, and 2, respectively; and IGL included 32, 20, and 8. Intergenic variants represented the largest class, with 4,166, 1,995, and 1,112 significant variants associated with IGHV, IGKV, and IGLV usage, respectively. Within IGH, IGHD usage was associated with 21 coding, 6 RSS, and 504 intergenic variants, whereas IGHJ usage was associated with 4 coding, 1 RSS, and 43 intergenic variants. Thus, genetic associations with segment usage span both the genes themselves and their surrounding non-coding sequence.

Repertoire usage also varied for IGHD and IGHJ ASCs represented in the corresponding RSS maps (Sup. Figure 6). We therefore asked whether D–J repertoire organization contained relationships between recombined segments that were not captured by their individual gene usage. Conditional usage was therefore calculated at the ASC level, first focusing on the 25 IGHD and six IGHJ ASCs retained for D–J analysis. For each D ASC, *P* (*J* | *D*) describes the relative distribution of J segments observed with that specific D, whereas *P* (*D* | *J*) describes the complementary distribution of D segments associated with a given J (Figure 6A–B).

The *P* (*J* | *D*) profiles differed markedly among IGHJ ASCs (Figure 6A). J4 and J6 showed strongly opposing D-usage profiles (*ρ* = −0.84), whereas J6 and J5 were highly concordant (*ρ* = 0.71), and J5 and J4 were strongly anticorrelated (*ρ* = −0.72). J1, J2, and J3 were more similar, with pairwise correlations ranging from *ρ* = 0.44 to 0.69. These patterns indicate that differences among J segments extend beyond their overall usage frequencies to the D backgrounds in which they occur.

The complementary *P* (*D* | *J*) analysis revealed local structure across the IGHD locus (Figure 6B). An upper concordant region included D5-24/D4-23 (*ρ* = 0.83), D4-23/D3-22 (*ρ* = 0.77), and D3-22/D2-21 (*ρ* = 0.83). A second concordant region lower in the locus contained neighboring profiles with correlations ranging from *ρ* = 0.66 to 0.77, including D4-11*/D3-10, D3-10/D3-9, D3-9/D1-7, D6-6/D3-3, and D3-3/D2-2*. These regions were separated by a discordant middle portion of the locus, with the strongest adjacent transition between D4-17 and D3-16 (*ρ* = −0.14). Conditional D usage therefore forms locally concordant regions separated by changes in D–J usage preference rather than following a continuous relationship with chromosomal position.

The same conditional framework was applied to IGH V–D relationships and to V–J relationships in IGK and IGL (Sup. Figure 8; Sup. Figure 9), showing that structured conditional segment usage is not restricted to IGH D–J rearrangements.

We next tested whether genetic variation was associated with conditional D–J usage. For each IGH variant and IGHJ ASC, D-specific enrichment values derived from *P* (*J* | *D*) were tested jointly across D backgrounds using a multivariate association test. Thus, rather than testing whether genotype changed the overall usage of a J segment, this analysis tested whether genotype was associated with its distribution across D backgrounds. Among all variant–J tests, *igh 37391* had the lowest association *p*-value and was selected for visualization in Figure 6C. The minimum association occurred for the J4 enrichment profile (Pillai = 0.737, *p* = 1.96 *×* 10^−16^).

The individual and conditional gene usage analyses of *igh 37391* produced markedly different patterns. Individual gene usage was significantly associated with genotype for 8 of 25 IGHD ASCs, whereas none of the six IGHJ ASCs passed the multiple-testing threshold. In contrast, 39 of 150 D–J combinations (26%) showed significant genotype-associated differences in D-specific enrichment derived from *P* (*J* | *D*). These associations were concentrated among J6 (13 combinations), J4 (11), and J3 (11), with four associations for J1 and none for J2 and J5. Thus, genotype was associated with the distribution of J usage across D backgrounds despite the absence of a detectable association with overall J usage.

The conditional associations were also not explained solely by changes in individual D usage. Of the 39 significant D–J enrichment associations, 27 (69%) involved D ASCs without a significant individual gene to genotype association. Because no J ASC was marginally significant, these 27 D–J associations were not detected by the individual gene level analysis of either component segment. For example, the individual usage of D3-3 did not have a guQTL that passed the multiple-testing threshold (*p* = 0.003), but it did show strong genetically associated conditional enrichment associations with J4 (*p* = 1.3 *×* 10^−9^) and J6 (*p* = 2 *×* 10^−8^). Conversely, the individual usage of D6-19 had the strongest guQTL association (*p* = 2 *×* 10^−16^) but only one significant conditional D–J association. These examples distinguish genetic effects on the abundance of individual segments from effects on the composition of their recombination partners.

Overall, conditional effects were observed across 20 of the 25 IGHD ASCs. Several D segments with significant conditional genotype associations, including D3-10, D3-9, and D3-3, also occurred within the lower concordant region identified in the *P* (*D* | *J*) analysis. Together, these results show that inherited variation can be associated with D–J repertoire composition independently of detectable changes in individual gene segment usage, revealing genetic effects on repertoire organization that are not captured by conventional guQTL analysis.

### Genotype similarity is associated with exact IGK CDR3 amino acid sharing

Extending the repertoire biases observed within individuals, we next asked whether germline variation is associated with repertoire features shared across individuals. We previously showed that *cis* genetic variants in heavy- and light-chain loci can influence gene usage patterns and CDR3 properties ^6,7^. Here, we examined this question at higher resolution by quantifying exact CDR3 amino acid (AA) sequence sharing across individuals. Shared CDR3 AA sequences were defined by exact sequence identity, without clustering or fuzzy matching. Exact CDR3 sharing was observed across IGH, IGK, and IGL, but was substantially more limited in IGH than in the light-chain repertoires (Figure 7A; Sup. Figure 10A). We therefore focused the main analysis on IGK.

To test whether CDR3 sharing was related to inferred genotype similarity, we compared pairwise IGK CDR3 AA overlap with pairwise overlap of PIgLET-inferred repertoire genotype sets (RGSs) (Figure 7B). CDR3 AA overlap increased across RGS-overlap bins, with median overlap coefficients of 0.481, 0.490, 0.496, and 0.501 from the lowest to the highest bin. Thus, individuals with more similar inferred IGK allele sets tended to share more exact IGK CDR3 AA sequences. In contrast, IGL showed little change in CDR3 AA overlap across increasing RGS-overlap bins despite substantial exact CDR3 sharing (Sup. Figure 10B), indicating that the relationship between genotype similarity and CDR3 overlap was not a general feature of light-chain repertoires.

We then examined shared IGK CDR3 AA sequences at the V-allele level (Figure 7C). V allele-specific shared CDR3 AA sequences were defined as exact CDR3 AA sequences observed in at least five individuals and assigned to a single IGK V allele. To focus on allele-associated sharing rather than broadly V-gene-associated sharing, V alleles observed in more than 60% of individuals were excluded. Shared CDR3 AA sequences were concentrated among a small set of V alleles, including IGKV1D-13*02, IGKV1-39/IGKV1D-39*01, IGKV2-29*02, and IGKV1-9*02. These sequences differed both in the number of individuals sharing them and in their median within-individual frequency. Additional V-allele-associated sharing in IGL is shown in Sup. Figure 10C.

Because IGKV1D-13 contributed prominently to the shared IGK CDR3 repertoire, we next examined genetic variation associated with its repertoire usage. A guQTL scan across the IGK locus identified a strong association with IGKV1D-13 usage, with *chr2 90219902* representing the lead variant (Figure 7D). IGKV1D-13 usage differed markedly across lead-variant genotypes, with progressively higher usage from 0/0 (*n* = 70), to 0/1 (*n* = 35), to 1/1 (*n* = 7) individuals (Figure 7E).

We then asked whether this genetically associated difference in IGKV1D-13 usage was reflected in the shared CDR3 repertoire. The within-individual frequency of the most widely shared IGKV1D-13-associated CDR3 AA sequence increased with IGKV1D-13 gene usage (Figure 7F). Individuals with genotypes associated with higher IGKV1D-13 usage also tended to show higher frequencies of this shared CDR3 sequence, linking inherited variation in gene usage to variation in a recurrent repertoire feature.

Together, these analyses show that inherited IGK variation is associated with exact CDR3 sharing at multiple levels. Individuals with more similar IGK repertoire genotypes share more exact CDR3 AA sequences, shared CDR3s are concentrated among a subset of V alleles, and genetic variation associated with IGKV1D-13 usage is also associated with the abundance of a recurrent IGKV1D-13-derived CDR3. These results indicate that inherited germline variation can contribute to public light-chain repertoire structure through both allele content and gene-usage differences.

## Discussion

Our results reveal multiple levels at which inherited immunoglobulin variation is associated with the organization of human antibody repertoires. Three principal findings emerge from the analyses presented in the paper. First, human IG diversity is substantially underrepresented in current reference sets and differs across ancestry groups and loci. Second, coding alleles are embedded within recurrent local sequence architectures: coding-region similarity is correlated with similarity in linked RSSs and leaders, although multiple coding alleles can share the same flanking sequence contexts. Third, inherited IG variation is associated with repertoire organization at levels not captured by marginal gene usage alone, including conditional D-J relationships and exact IGK CDR3 amino acid sharing across individuals. Together, these findings support a genomic-context-aware model in which IG coding alleles are interpreted in the context of linked gene-adjacent and locus-level variation.

The Human Unified Set of Alleles (HUSA) expands the baseline reference from 401 to 1,447 alleles and provides genomic and repertoire evidence for variation across all three IG loci. Many non-baseline alleles were observed in multiple individuals, indicating that incomplete reference representation is not limited to rare, sample-specific sequences. Within the sampled cohorts, individuals of African ancestry carried the largest number of observed alleles and showed the lowest within-ancestry allele-set overlap across all three loci, consistent with earlier reports of substantial IG diversity in populations of African ancestry ^8,39,40^. IGH also showed lower within-ancestry over-lap than IGK across all population groups, consistent with greater inter-individual diversity in the sampled IGH data. This observation complements previous single-genome comparisons reporting enrichment of genetic variation within IGH relative to the genome average and the light-chain loci ^41^. These comparisons should nevertheless be interpreted in light of differences in locus structure, annotation complexity, reference completeness, and retained sample counts. More broadly, the results emphasize that incomplete and ancestry-imbalanced germline references can affect repertoire annotation, genotype inference, and comparisons among populations.

Beyond expanding the allele catalog, HUSA reveals that coding and gene-adjacent variation are non-randomly organized. Pairwise coding-region distance was positively correlated with linked RSS and leader distance across IGHV, IGKV, and IGLV, and coding-RSS concordance persisted within sufficiently represented V-gene subgroups. At the same time, flanking-sequence diversity was compressed relative to coding diversity, with multiple coding alleles and, in some cases, multiple genes sharing recurrent RSS or leader sequences. These complementary observations indicate that IG alleles are embedded within partially conserved local sequence architectures rather than carrying unique flanking contexts. In this view, the relevant inherited unit is not only a coding variant within a gene, but a local genomic context that includes linked RSS and leader variation.

Matched genomic and repertoire data further expanded the set of RSS and leader sequence classes linked to alleles recovered from productive repertoires. This advances previous descriptions of gene-adjacent sequences that were based on more restricted genomic sampling and provides population-scale evidence for a broader range of coding-RSS-leader combinations. However, recovery of a linked allele from productive AIRR-seq does not establish the functional contribution of its RSS or leader sequence independently of coding variation, genomic position, cellular selection, or other linked regulatory features. Direct recombination and expression assays will therefore be required to determine which of these sequence variants alter repertoire formation.

RSS organization also differed among sequence components. Heptamer and nonamer motifs were comparatively conserved, consistent with their roles in RAG-mediated recognition, cleavage, and enforcement of the 12/23 rule, whereas spacer sequences accounted for a larger proportion of complete-RSS divergence ^42–45^. Analyses restricted to consensus heptamer and nonamer motifs may therefore miss a substantial component of inherited RSS variation. Although previous experimental work indicates that spacer sequence can influence recombination efficiency ^45^, the present analysis does not establish whether the spacer variants observed here alter recombination directly. Testing selected variants in controlled recombination assays will be needed to distinguish functional effects from sequence divergence arising through linkage or shared evolutionary history.

The observed concordance between coding and gene-adjacent sequences should not be interpreted as a formal phylogeny or as evidence that coding regions, RSSs, and leaders share identical evolutionary histories. Rather, the pattern is consistent with duplication and diversification of local sequence blocks, followed by differential constraint, gene conversion, recombination, or other remodeling processes affecting their individual components ^5,46,47^. Complete haplotype-resolved comparisons across IG loci will be needed to distinguish among these possibilities and to determine whether recurrent flanking contexts reflect shared ancestry, sequence exchange, or convergent constraint. Nevertheless, the present results argue that coding alleles should not be treated as entirely independent of their linked genomic contexts.

The conditional segment-usage analysis extends conventional gene-usage QTL approaches by testing dependencies between rearranged segments rather than treating each segment independently. Conditional D-J profiles showed structured relationships across the IGH locus, consistent with previous evidence for positional biases in D-J pairing and with models in which recombination is shaped by locus architecture, chromatin context, and segment accessibility ^27–29,48^. Importantly, the representative genetic variant was associated with D-specific J enrichment even though no J allele similarity cluster showed a significant marginal association with genotype. Moreover, 27 of the 39 significant conditional D-J associations involved D clusters without significant marginal genotype associations. Inherited effects on repertoire organization can therefore be visible in recombination-partner distributions even when they are not detected through the abundance of either component segment alone. This distinction suggests that part of the inherited signal in repertoire organization resides in relationships among rearranged segments rather than in marginal segment usage alone. Probabilistic models of repertoire formation may therefore benefit from representing genotype-associated segment-pair dependencies in addition to marginal V, D, and J usage.

These conditional associations should not yet be interpreted as evidence that the tested variants directly control D-J pairing. Genetic markers across the IG loci may capture combinations of linked coding, regulatory, structural, and positional features. In addition, the conditional analysis was performed in a modest-sized cohort, did not include additional ancestry or technical covariates, and used cell-level tests as follow-up analyses within a selected significant multivariate profile. The individual variant-pair relationships will therefore require replication in larger independent cohorts and, ultimately, experimental or haplotype-resolved analyses capable of identifying the causal genomic features.

The IGK sharing analysis provides complementary evidence that inherited variation is associated with repertoire features shared across individuals. Pairs of individuals with more similar inferred IGK allele sets showed an increase in exact CDR3 amino acid overlap, although a corresponding relationship was not evident for IGL. In addition, a variant strongly associated with IGKV1D-13 usage marked individuals with different frequencies of a recurrent IGKV1D-13-associated CDR3. These observations are consistent with inherited allele content and gene-usage biases contributing to the probability of generating or retaining recurrent light-chain sequences, extending previous findings that IG variants are associated with gene usage, allele representation, and broader CDR3 properties ^6,7^.

The CDR3-sharing results also require careful interpretation. They do not establish direct mediation by the lead IGKV1D-13-associated variant, and exact identity of an unpaired light-chain CDR3 does not establish identity of the complete antibody or shared antigen specificity. Repertoire depth, clonal structure, developmental selection, environmental exposure, and convergent recombination may also contribute to exact sequence sharing. Furthermore, the pairwise repertoire comparisons are not statistically independent because each individual contributes to multiple pairs. Replication using independent cohorts, paired heavy-light sequencing, and models that account for repertoire depth, ancestry, clonal structure, and non-independence among subject pairs will be needed to determine the magnitude and functional significance of this relationship.

Together, the results support a multilayer model of baseline repertoire formation. Inherited allele content defines the available coding substrates, while linked RSSs, leaders, genomic position, structural variation, and broader chromatin context may contribute to recombination accessibility and segment usage ^26–29,49,50^. Genetic effects may also be expressed through dependencies between rearranged segments and through the recurrence of specific CDR3 sequences. The present analyses do not assign causal weights to these layers or distinguish the effects of individual linked features. Instead, they identify relationships that should be represented explicitly in future haplotype-aware statistical models and tested through experimental perturbation.

Several limitations should be considered. First, portions of the genomic datasets, including samples from the 1KGP and HPRC, were derived from lymphoblastoid cell lines (LCLs). LCL-derived material can contain somatic rearrangements and mutations that complicate assembly, annotation, and genotype inference across IG loci ^5,19,47,51,52^. Although HiFi long reads and stringent contig-level and read-level support requirements were used to mitigate these effects, V(D)J rearrangement can result in partial haplotype loss and biased genotype or allele-frequency estimates unless explicitly modeled ^51^. The present analysis therefore emphasizes allele observations across samples and populations rather than population-frequency estimates that may be distorted by these artifacts.

Second, matched genomic and repertoire analyses were restricted to 172 individuals, limiting power for ancestry-stratified and multivariable association analyses. Third, recovery from productive AIRR-seq reflects the combined effects of recombination, transcription, cellular selection, repertoire sampling, and inference thresholds; it is not a direct assay of RSS or leader function. Fourth, the association analyses do not distinguish causal variants from linked markers, and the absence of additional genetic, ancestry, or technical covariates may permit residual confounding. Finally, the present analyses do not fully represent structural variation, chromosomal position, three-dimensional locus organization, or complete haplotypic context, each of which may contribute to the observed relationships. Independent non-LCL cohorts, complete haplotype-resolved assemblies, and direct functional validation will therefore be important next steps.

In summary, this study supports a shift from gene-centric descriptions of IG variation toward architecture-aware models in which coding alleles are considered together with linked RSSs, leaders, and locus-level genomic variation. By connecting population-scale allele diversity with expressed allele recovery, local sequence organization, conditional segment usage, and exact CDR3 sharing, the study identifies multiple levels at which inherited IG variation is associated with repertoire structure. HUSA provides an evidence-linked, long-read-based foundation for testing these relationships in larger and more diverse cohorts, improving germline gene and allele annotation, and developing genotype-aware models of immune receptor generation. Its open-access framework will also support the incorporation of future long-read datasets, including data generated from non-LCL materials and complete haplotype-resolved genomes. As increasingly complete haplotype-resolved IG genomes become available, integrating coding, regulatory, structural, and repertoire variation should enable increasingly accurate models of the inherited genomic architectures that shape human antibody repertoires. Establishing which linked features act causally will require independent matched genomic-repertoire cohorts and direct functional assays of candidate regulatory variants.

## Methods

### Data Sources, Sample Collection, and Sequencing

This study integrated three data sources for population-scale analysis of human IG germline variation across the IGH, IGK, and IGL loci. DS1 consisted of 1KGP samples with targeted long-read genomic data. DS2 consisted of HPRC samples with long-read whole-genome data. DS3 leveraged previously published data generated from PBMCs collected from healthy donors (procured from STEMCELL Technologies) with matched targeted long-read genomic data and AIRR-seq repertoires. The acquisition of de-identified DNA and PBMCs was approved by the Institutional Review Board of the University of Louisville. Dataset labels, data types, retained locus-level sample counts, and ancestry or population groups are summarized in Table 1.

For DS1, genomic DNA isolated from LCLs was procured from the Coriell Institute for Medical Research (Camden, NJ) for individuals previously collected as part of the 1KGP. Genomic DNA was processed using our previously published targeted long-read IGH, IGK, and IGL sequencing protocol ^7,8,47^. This approach allows for sample-level targeted pull-down of long DNA fragments representing the IG loci, which can be used to generate IG loci-enriched sequencing libraries that can be multiplexed and sequenced on any Pacific Biosciences platform. Libraries generated for this study were sequenced either on Sequel I, II, or Revio systems. The same targeted sequencing approach was used to generate data for the DS3 cohort, including locus-wide HiFi reads for IGH, IGK and IGL. Data for the IGK and IGL loci were previously published alongside their matched IgK and IgL AIRR-seq data ^7^. For this paper, we additionally analyzed and reported on the IGH locus data generated for these samples, as well as newly generated matched AIRR-seq libraries for the IgM repertoire. Briefly, to construct AIRR-seq libraries, RNA was isolated from ~3–5 million PBMCs per donor via the Qiagen AllPrep DNA/RNA Kit ^7^. AIRR-seq libraries were subsequently generated using a 5’ RACE approach. IgM 5’RACE libraries were prepared using the SMARTer Human BCR Profiling Kit (Takara Bio, San Jose, CA, USA). Library quality and concentration for indexed IgM libraries were evaluated using the 2100 Bioanalyzer High Sensitivity DNA Assay Kit (Agilent, Santa Clara, CA, USA) and the Qubit 3.0 Fluorometer dsDNA High Sensitivity Assay Kit (Life Technologies, Carlsbad, CA, USA), respectively. Pooled libraries (10 nM) were sequenced on the Illumina NextSeq platforms using 300 bp paired-end reads. No matched AIRR-seq data were generated for DS1 or DS2, and those samples were excluded from repertoire-based analyses. Cohort metadata were harmonized into AFR, AMR, EAS, EUR, SAS, and Mixed categories.

### Baseline Reference Set

The baseline reference set was used as the seed for genomic annotation, allele harmonization, and classification of baseline versus non-baseline alleles. The baseline set was constructed from the latest version of the OGRDB human IG reference available at the time of analysis, supplemented with additional IMGT alleles not represented in OGRDB. The set included productive V, D, and J alleles from the IGH, IGK, and IGL loci. Retained alleles matching this baseline set were classified as baseline alleles, whereas retained productive alleles absent from the baseline set were classified as non-baseline alleles. The same baseline reference set was used across all data sources. The relationship among the baseline reference, HUSA, and IMGT was evaluated per gene segment by comparing retained allele coding-region sequences using exact sequence identity. The IMGT comparison set was restricted to functional and ORF alleles of genes represented in HUSA.

### Genome Assembly and WASP Annotation

IGH, IGK, and IGL locus-specific PacBio HiFi reads were used to generate haplotype-phased de novo assemblies using the WASP ^37^ framework, which employs Hifiasm (v0.19) with default parameters to produce primary phased contigs. Because many samples had high IG-locus read depth, reads were preprocessed before assembly to reduce LCL-related rearrangement artifacts and to normalize coverage across samples. As we and others have noted, genomic reads generated from LCL-derived DNA can contain V(D)J recombination events, which can result in erroneous assembly contigs and variant calls ^5,51,52^. To mitigate these effects, we first filtered reads containing evidence of V(D)J rearrangement using a custom Python script. Specifically, the script excluded reads exhibiting region-specific multi-mappability, defined as either primary or supplementary alignments mapping to both V and J gene regions. The filtered read set was then downsampled to 50*×* coverage using Rasusa ^53,54^, while preserving the original read-length distribution, and used for downstream genome assembly.

The resulting phased assemblies were concatenated to remove redundancy and aligned to a custom reference assembly using Minimap2 (v2.26) with the ‘-x asm20’ preset. Alignment was performed iteratively, beginning with initial placement of IG-specific contigs, followed by resolution of partial alignments through realignment of unmapped soft-clipped regions. During each iteration, soft-clipped segments were extracted and realigned to refine primary alignments. The final phased assembly contig alignments were then processed through the WASP annotation steps to curate variants and gene/allele annotations. Genomic allele calls were annotated against the baseline IG reference set used as the seed for downstream allele naming, harmonization, and clustering. WASP-based annotation was applied across loci and was also used to extract RSS and leader sequences from allele-flanking genomic regions for downstream RSS, leader, and usage analyses.

### Genomic Allele Retention and Germline Set Construction

GGSs were constructed from productive genomic allele calls after allele-level quality control. Baseline alleles were retained if the sequence accuracy metric was 100%. Non-baseline alleles were retained if they had a sequence accuracy metric of 100% and support from at least 10 fully spanning reads with a 100% match. Non-baseline alleles that did not meet the read-support criterion in a given subject were also retained if the same allele was observed in another subject from the same genomic dataset and met the full qualifying criteria in that subject.

Pseudogene allele calls were removed before GGS construction. Allele calls with annotation notes indicating incomplete coding-region annotation, missing RSS or leader features, or sequence truncation were excluded before GGS construction. These filters were applied before constructing locus-specific GGSs.

DS1 and DS2 were processed as genomic-only resources and used as additional allele support sources for HUSA integration. Because DS1 and DS2 did not have matched repertoire data, no repertoire-depth filtering or RGS inference was performed for these samples.

### DS3 Matched-Sample Inclusion Criteria

For DS3, an additional matched-sample inclusion filter was applied after genomic allele retention. This filter was used for downstream analyses that integrated genomic alleles with repertoire-derived measurements, including expressed allele recovery, RSS and leader analyses, gene-usage analyses, conditional V(D)J usage, and CDR3 sharing. The purpose of this filter was not to define whether an individual allele call was acceptable, but to ensure that each retained subject-locus had broadly supported genomic characterization before being used in matched genomic-repertoire analyses.

For each DS3 subject and locus, the subject-locus was retained if its repertoire had a minimum depth of 1,000 sequences for the corresponding locus. From this point onward, DS3 analyses used only subject-loci passing both the genomic-support and repertoire-depth criteria, unless otherwise specified. This subject-level filter provided an additional layer of support beyond allele-level retention criteria for analyses that depended on matching genomic and repertoire evidence within the same individual.

### AIRR-seq Processing, Novel Allele Inference, and RGS Inference

Raw AIRR-seq reads from DS3 were preprocessed with nf-core/airrflow using the takara smartseq umi bcr library preparation profile. AIRRflow was used only for read preprocessing and generation of processed repertoire sequence files, not for downstream V(D)J annotation, genotype-aware alignment, allele filtering, or RGS inference.

For each DS3 subject and locus, processed AIRR-seq sequences were annotated against the subject-specific genomic reference constructed from the corresponding GGS. IgBLAST-based V(D)J alignment was performed separately for IgM, IgK, and IgL libraries. Heavy-chain IgM reads were annotated against IGHV, IGHD, and IGHJ references, whereas IgK and IgL reads were annotated against the corresponding V and J references.

Novel V alleles were inferred using TIgGER from sequences aligned to the personalized references. Candidate alleles were used to update the subject-specific reference set, after which sequences were realigned. For heavy chains, clones were assigned using Change-O, and a minimally mutated representative sequence was selected for each clone. Downstream genotype inference used sequences with zero nucleotide mutations in the V-gene region.

To infer RGSs, allele-specific inclusion thresholds were optimized using PIgLET, which combines allele similarity clustering with a matched genomic-training strategy for genotype inference ^20,38^. Threshold optimization compared AIRR-seq-inferred alleles to matched GGS calls in a training cohort. Repertoire rows were retained for threshold training when the subject-locus depth was at least 1,000 sequences. V- and D-gene calls with multiple assignments were excluded from allele counting, whereas J-gene calls with multiple assignments were split across assigned alleles with fractional weights.

Eligible repertoires were split 50:50 into training and held-out sets using a fixed random seed, with stratification by V-segment depth quantiles within each locus. For each allele, a grid of candidate frequency thresholds was evaluated against the matched genomic truth set, and the lowest threshold maximizing the mean per-sample F1 score was selected. This optimization used a Z-score cutoff of 0 during threshold selection. The optimized allele-specific thresholds were written as a reusable PIgLET threshold table.

Following threshold optimization, RGSs were inferred using PIgLET. For each subject and locus, an allele was included in the RGS if its observed repertoire frequency exceeded the optimized allele-specific threshold and its Z-score was greater than 1. This combined frequency and confidence filter was used to define expressed alleles for downstream analyses.

### HUSA Construction and Allele Similarity Clustering

The Human Unified Set of Alleles (HUSA) was constructed by integrating the baseline reference set, DS1 and DS2 genomic-only support, and DS3 GGSs and RGSs. Genomic records from all resources were harmonized to a common sequence- and allele-based schema, after which repertoire support was merged onto matching genomic records. Repertoire-derived alleles were included only when they met RGS inference criteria.

Identical coding sequences across individuals and data sources were collapsed, and allele metadata were annotated with source-specific support. DS1 genomic support, DS2 genomic support, DS3 genomic support, and DS3 repertoire support were retained as separate evidence fields. When the same allele was observed in both genomic and repertoire data, the genomic allele record was used as the primary sequence record, and repertoire evidence was retained as support metadata.

Allele similarity clustering was performed across V, D, and J segments using the method implemented in PIgLET and described previously ^20,38^. Pairwise dissimilarity matrices were constructed from Hamming or Levenshtein distances, depending on whether sequences could be compared as aligned strings. Dissimilarity matrices were transformed into similarity networks and clustered with the Leiden algorithm using a Constant Potts Model objective function.

### Allele Prevalence, Ancestry Overlap, and Expressed Allele Analyses

Allele prevalence in genomic datasets was calculated as the number of retained individuals carrying each HUSA allele within each locus. Alleles were ranked by carrier count for rank-distribution plots. Ancestry-stratified allele overlap was calculated from HUSA alleles with genomic support and visualized with UpSet plots by locus. Within-ancestry overlap coefficients were calculated between pairs of individuals as the size of the shared genomic allele set divided by the size of the smaller genomic allele set.

Expressed allele prevalence was calculated from DS3 RGSs as the number of subjects whose RGS contained each allele. Expressed allele rank distributions were generated separately by gene segment, with alleles labeled as baseline or non-baseline according to the baseline reference set.

### RSS and Leader Sequence Analyses

RSS and leader sequences were extracted by WASP from allele-flanking genomic regions. RSSs were represented as heptamer-spacer-nonamer triplets, and spacer lengths were recorded for each allele-RSS pair. Leader exon 1 and exon 2 sequences were extracted from the corresponding upstream regions by the same annotation workflow.

Consensus categories were assigned separately for RSS and leader sequences. For RSSs, variants were classified as “Both”, “Either”, or “None” according to whether both, one, or neither of the heptamer and nonamer matched the gene-segment-specific consensus motifs. For leaders, variants were classified analogously according to whether both, one, or neither of leader exon 1 and exon 2 matched the corresponding gene-segment-specific consensus sequences. RSS analyses used unique allele-RSS triplets from the DS3 genomic export used for matched RSS and leader analyses. Heptamer and nonamer sequence logos were weighted by the number of unique allele-RSS sequence pairs. RSS variants were compared with IMGT by matching full heptamer-spacer-nonamer triplets within each gene segment. HUSA-only, IMGT-only, and shared RSS variants were summarized according to source and whether the linked allele was observed in repertoire data.

Leader sequences from HUSA genomic records were compared with IMGT leader sequences by V-gene locus. Unique leader sequences were summarized by source overlap, carrier count, and whether the linked HUSA allele was observed in expressed repertoire data.

Coding and non-coding sequence contexts were evaluated from unique allele-RSS and allele-leader combinations. For each V-gene locus, we calculated the numbers of unique coding alleles, complete RSS sequences, heptamernonamer combinations, spacer sequences, and full-length leader sequences. Sharing of RSS and leader sequences across coding groups was quantified by the number of genes associated with each unique sequence. IGHD 5′ and 3′ RSSs were analyzed separately.

Relative repertoire usage was calculated from matched genotype-aware repertoire annotations after collapsing allele calls to ASCs. For each subject and locus, relative usage was calculated as the fraction of retained repertoire sequences assigned to each ASC.

### RSS Conservation and Coding Noncoding Sequence-Distance Concordance

For RSS conservation analyses, nucleotide-distance matrices were computed separately for RSSs with 23-bp and 12-bp spacers. Within each spacer class, distances were calculated for the complete heptamer-spacer-nonamer RSS, the combined heptamer and nonamer motifs, and the spacer alone. Hamming distance was used when all sequences in a comparison had the same length, and Levenshtein distance otherwise. Distances were normalized to the range [0, 1] by dividing Hamming distances by sequence length and Levenshtein distances by the maximum observed distance within the corresponding sequence set.

To quantify the contribution of individual RSS components to complete-RSS divergence, Mantel correlations were calculated between the complete-RSS distance matrix and the corresponding spacer-only and heptamer-plus-nonamer distance matrices. Nucleotide diversity was also calculated separately for the heptamer, spacer, and nonamer. For each component and spacer class, per-site nucleotide diversity was defined as the mean pairwise Hamming distance among component sequences represented by the set of unique complete RSSs, divided by the component length. This measure corresponds to the mean proportion of nucleotide positions that differ between sequence pairs and ranges from 0 to 1. As a complementary measure of sequence variability, the number of polymorphic nucleotide positions within each component was also calculated.

To assess concordance between coding-region and linked non-coding sequence variation, we compared pairwise sequence distances within each IG locus. For each allele, the coding-region sequence was paired with the corresponding RSS or full-length leader sequence obtained from the DS3 genomic RSS and leader annotations. Only alleles for which both the coding sequence and the corresponding linked sequence element were available were included.

Pairwise coding-region and leader distances were calculated using Levenshtein distance. RSS distances were calculated using Hamming distance when all aligned RSS sequences had the same length and Levenshtein distance otherwise. Concordance between coding-region and linked sequence distances was assessed using Mantel tests with Spearman correlation and 4,000 permutations of allele labels. One-sided permutation *p* values tested whether the observed correlation was greater than expected under the permutation null distribution.

Tests were performed for complete RSS and full-length leader sequences in IGHV, IGKV, and IGLV, and for RSS sequences in IGHJ, IGKJ, IGLJ, and the 5′ and 3′ flanks of IGHD. Mantel test *p* values were corrected together using the Benjamini–Hochberg false discovery rate procedure.

To determine whether coding–RSS concordance was explained primarily by divergence between V-gene subgroups, coding–RSS Mantel analyses were additionally performed within individual V subgroups. Only subgroups containing at least six alleles with paired coding-region and RSS sequences were included.

### Gene-Usage QTL Analyses

Gene-usage quantitative trait locus (guQTL) analyses were performed in DS3 using matched IG-locus variant genotypes and genotype-aware repertoire annotations. For each retained V, D, or J ASC, relative repertoire usage was calculated for each individual from the corresponding locus-specific repertoire.

Relative usage was defined as the proportion of sequences assigned to an ASC among all sequences assigned to the corresponding segment class within that locus. To obtain finite values when an ASC had zero counts, a pseudocount of 0.5 was added to each ASC count, with the denominator increased by 0.5 multiplied by the number of ASCs in the corresponding segment class. The resulting proportions were logit-transformed before association testing.

For each segment class, eligible genetic variants were tested individually for association with relative ASC usage. Variants were encoded additively as alternative-allele dosage (0*/*0, 0*/*1, or 1*/*1). For each variant–ASC pair, logit-transformed usage was regressed on genotype dosage using univariate linear regression. Association *p* values were obtained from a two-sided *t*-test of the genotype regression coefficient. No additional covariates were included. IGK and IGL guQTL analyses followed the previously reported framework ^7^, and the same framework was applied to IGH in the present study.

Variants were retained if their minor-allele frequency was at least 0.05. Missing genotypes were not imputed. Variants sharing the same genotype-missingness pattern were analyzed using the corresponding complete-case subject set, and at least 60 individuals with non-missing genotype and repertoire measurements were required for a variant to be tested.

Multiple testing was controlled separately within each locus using a Bonferroni threshold of *α/m*, where *α* = 0.05 and *m* was the number of independent variants remaining after variants in complete linkage disequilibrium were collapsed. A variant–ASC association was considered significant when its *p* value was below the corresponding locus-specific threshold. Independent lead variants were subsequently identified by linkage-disequilibrium clumping at *r*^2^ *≥* 0.80. Lead variants for which the smallest genotype class contained at least five individuals were retained for genotype-stratified follow-up analyses.

Significant variants were annotated according to their genomic position relative to IG coding regions, leader sequences, RSSs, and intergenic regions. Significant guQTL variants were summarized separately by locus, segment class, and genomic annotation.

### Conditional V(D)J Usage Analyses

Conditional V(D)J usage analyses used productive DS3 repertoire sequences after genotype-aware alignment and allele clustering, with alleles mapped to ASCs before counting. For IGH, the conditional probabilities *P* (*J* | *D*), *P* (*D* | *J*), *P* (*D* | *V*), and *P* (*V* | *D*) were calculated from each subject’s rearrangements. For IGK and IGL, the corresponding V–J probabilities *P* (*J* | *V*) and *P* (*V* | *J*) were calculated. Conditional probabilities were summarized across subjects as the median for each segment pair.

Subjects were excluded from IGHD and IGHJ usage analyses when genomic processing did not retain a call for a gene present in that subject. Such a gene is absent from the subject-specific reference, so its rearrangements are annotated against the remaining genes rather than discarded. A segment was retained only if it was observed at least 10 times in each of at least 100 subjects, and per-subject events for a retained segment required at least 10 supporting sequences. Unobserved ASC pairs were left as missing and excluded from correlation analyses using pairwise complete observations; no pseudocount was added.

Pairwise Spearman correlations were computed between conditional-usage profiles. IGHD ASCs were ordered by seriation followed by the genomic position of the first gene in each cluster. IGH V–D and light-chain V–J conditional-usage profiles were analyzed using the same framework.

To test whether inherited genetic variation was associated with conditional D–J usage, a D-specific enrichment value was calculated for each IGHD–IGHJ ASC pair and individual. For subject *s* and pair (*d, j*), let *n*_*sdj*_ denote the rearrangement count, *n*_*sd·*_ and *n*_*s·j*_ the corresponding IGHD and IGHJ marginal counts, and *N*_*s*_ the subject’s total number of rearrangements. Under independence of D and J usage, the expected count was

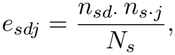

and enrichment was defined as

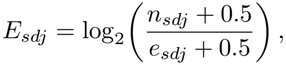

where a pseudocount of 0.5 was added to maintain finite values for zero c ounts. In the absence of the pseudocount, this quantity is equivalent to log_2_ *P* (*J* | *D*) − log_2_ *P* (*J*), and therefore measures D-specific d eviation f rom t he J usage expected under independent segment usage.

For each IGHJ ASC anchor, the response was the vector of D-specific enrichment values across retained IGHD ASCs. Variants with a minor-allele frequency of at least 0.05 were eligible for testing. For a given anchor, IGHD responses were retained according to the required subject-level data-availability criterion, and the anchor was tested only when at least two IGHD responses and at least 60 subjects with complete data remained. Each retained anchor was tested against every eligible variant using multivariate analysis of variance, with genotype encoded additively as alternative-allele dosage (0*/*0, 0*/*1, or 1*/*1) and Pillai’s trace used as the multivariate test statistic.

Multiple testing was controlled using the same locus-specific variant space as in the marginal gene-usage analysis. A Bonferroni threshold of *α/m* was applied, where *α* = 0.05 and *m* was the number of variants remaining after variants in complete linkage disequilibrium were collapsed. A variant–IGHJ association was considered significant when its multivariate *p* value was below this threshold. Genotype-class counts were calculated for each tested anchor, and associations for which the smallest genotype class contained at least five individuals were retained for genotype-stratified follow-up analyses.

For each variant IGHJ combination passing the multivariate significance threshold, individual IGHD–IGHJ cells were evaluated using univariate linear regression of the corresponding per-subject enrichment value on genotype dosage. A cell was considered significant when the anchoring multivariate association passed the Bonferroni threshold and the cell-level genotype-slope *p* value was below 0.05. Cell-level tests were performed only within significant multivariate profiles and were therefore interpreted as conditional follow-up tests rather than as independent tests across the complete IGHD *×* IGHJ matrix.

The variant *igh 37391* was selected for visualization because it produced the lowest *p* value across all tested IGH variant–IGHJ multivariate associations. Marginal genotype associations with IGHD and IGHJ usage were evaluated using the corresponding gene-usage QTL framework, allowing marginal and conditional genotype effects to be compared for the same variant.

### CDR3 Amino Acid Sharing Analyses

CDR3 amino acid sharing analyses used productive DS3 repertoire sequences passing repertoire filters. Exact CDR3 amino acid sequences were used without clustering or sequence-similarity matching. A CDR3 amino acid sequence was classified as shared when the exact amino acid sequence was observed in more than one individual. Complementary cumulative distribution functions summarized the number of individuals sharing each exact CDR3 amino acid sequence for IGH, IGK, and IGL.

For pairwise IGK and IGL repertoire analyses, CDR3 amino acid overlap was calculated between pairs of individuals as the size of the shared exact CDR3 amino acid set divided by the size of the smaller CDR3 amino acid set. Individual RGS overlap was calculated analogously from PIgLET-inferred repertoire genotype sets. Subject pairs were grouped into four bins using quartile cut points from the empirical RGS-overlap distribution. When repeated quantile cut points prevented four distinct intervals, four equal-width intervals over the observed range were used instead. Within each RGS-overlap bin, pairwise CDR3 amino acid overlap was summarized by its median and interquartile range. The relationship between RGS overlap and CDR3 overlap was evaluated descriptively across these bins.

V allele-specific shared CDR3 amino acid sequences were defined as exact CDR3 amino acid sequences observed in at least five individuals and assigned to a single V allele. V alleles observed in more than 60% of individuals in the corresponding locus were excluded from this analysis. For each retained CDR3 amino acid sequence, the number of individuals sharing the sequence and its median within-individual frequency were calculated.

For IGK, IGKV1D-13-associated shared CDR3 sequences were examined together with genetic variation associated with IGKV1D-13 repertoire usage. The lead guQTL variant was defined as the variant with the strongest association with IGKV1D-13 usage in the locus-wide gene-usage scan. The most widely shared IGKV1D-13-associated CDR3 amino acid sequence was selected for comparison with IGKV1D-13 gene usage. For each individual, IGKV1D-13 usage and lead-CDR3 usage were expressed as fractions of the total IGK repertoire. Lead-CDR3 usage was regressed on IGKV1D-13 usage with the intercept constrained to zero, providing an estimate of their proportional relationship. Individuals in whom the lead CDR3 was not observed were retained with a CDR3 usage of zero.

### Declaration of generative AI and AI-assisted technologies in the manuscript preparation process

During the preparation of this work, the authors used Claude Code to assist in drafting initial versions of scripts used for data processing, and ChatGPT for language-editing suggestions. The authors reviewed, edited, and validated the AI-assisted outputs as appropriate. No AI-assisted technology was used to generate the scientific conclusions. The authors take full responsibility for the content of the published article.

## Code availability

All analysis code and the per-figure source tables are available at https://github.com/ayeletperes/husa.

## Ethics, Consent to Participate, and Consent to Publish declarations

**Ethics approval:** The acquisition and use of de-identified human genomic DNA and peripheral blood mononuclear cells (PBMCs) for this study were approved by the Institutional Review Board (IRB) of the University of Louisville. The study also included analyses of previously published and publicly available genomic datasets. **Consent to participate:** Informed consent was obtained from the original donors. **Consent to publish:** Not applicable. The manuscript does not contain identifiable individual participant information.

## Ethics declaration

The study was approved by the Institutional Review Board of the University of Louisville. Human-derived samples and associated data were analyzed in de-identified form.

## Author Contribution

A.P., U.J., C.T.W., and G.Y. conceived and designed the study. M.L.S. and C.T.W. coordinated sample collection, sequencing, and genomic data generation.

U.J., O.L.R., Z.M.V., E.E., W.S.G., K.S., B.C., S.S., C.B., and C.M. contributed to data generation, processing, genomic assembly, annotation, and quality control.

A.P. and U.J. led the computational analyses and the integration of genomic and antibody repertoire data, with contributions from O.L.R., Z.M.V., E.E., and W.D.L.

A.P., U.J., C.T.W., and G.Y. interpreted the results and developed the scientific framework for the manuscript.

A.P. and U.J. prepared the figures and drafted the manuscript, with input from C.T.W. and G.Y. M.L.S., C.T.W., and G.Y. supervised the research.

All authors contributed to revising the manuscript and approved the final version.

## Acknowledgements

The study was supported by the National Institutes of Health [grant numbers R24 AI138963, and U24 AI177622].

## References

[1] K.M. Murphy and C. Weaver. Janeway’s Immunobiology: Tenth International Student Edition with Registration Card. Titolo collana. W.W. Norton, 2022.

[2] Valerie H Odegard and David G Schatz. Targeting of somatic hypermutation. Nature Reviews Immunology, 6(8):573–583, 2006.

[3] Bryan Briney, Anne Inderbitzin, Collin Joyce, and Dennis R Burton. Commonality despite exceptional diversity in the baseline human antibody repertoire. Nature, page 1, 2019.

[4] María Ruiz Ortega, Natanael Spisak, Thierry Mora, and Aleksandra M Walczak. Modeling and predicting the overlap of b-and t-cell receptor repertoires in healthy and sars-cov-2 infected individuals. PLoS Genetics, 19(2):e1010652, 2023.

[5] Corey T Watson, Karyn M Steinberg, John Huddleston, Rene L Warren, Maika Malig, Jacqueline Schein, A Jeremy Willsey, Jeffrey B Joy, Jamie K Scott, Tina A Graves, Richard K Wilson, Robert A Holt, Evan E Eichler, and Felix Breden. Complete haplotype sequence of the human immunoglobulin heavy-chain variable, diversity, and joining genes and characterization of allelic and copy-number variation. The American Journal of Human Genetics, 92(4):530–546, 2013.

[6] Oscar L Rodriguez, Yana Safonova, Catherine A Silver, Kaitlyn Shields, William S Gibson, Justin T Kos, David Tieri, Hanzhong Ke, Katherine JL Jackson, Scott D Boyd, et al. Genetic variation in the immunoglobulin heavy chain locus shapes the human antibody repertoire. Nature communications, 14(1):4419, 2023.

[7] Eric Engelbrecht, Oscar L Rodriguez, William Lees, Zach Vanwinkle, Kaitlyn Shields, Steven Schultze, William S Gibson, David R Smith, Uddalok Jana, Swati Saha, et al. Germline polymorphisms in the immunoglobulin kappa and lambda loci underpinning antibody light chain repertoire variability. Nature Communications, 2025.

[8] William S Gibson, Oscar L Rodriguez, Kaitlyn Shields, Catherine A Silver, Abdullah Dorgham, Matthew Emery, Gintaras Deikus, Robert Sebra, Evan E Eichler, Ali Bashir, et al. Characterization of the immunoglobulin lambda chain locus from diverse populations reveals extensive genetic variation. Genes & Immunity, 24(1):21–31, 2023.

[9] Moriah Gidoni, Omri Snir, Ayelet Peres, Pazit Polak, Ida Lindeman, Ivana Mikocziova, Vikas Kumar Sarna, Knut EA Lundin, Christopher Clouser, Francois Vigneault, et al. Mosaic deletion patterns of the human antibody heavy chain gene locus shown by bayesian haplotyping. Nature communications, 10(1):1–14, 2019.

[10] Andrew M Collins, Gur Yaari, Adrian J Shepherd, William Lees, and Corey T Watson. Germline immunoglobulin genes: disease susceptibility genes hidden in plain sight? Current Opinion in Systems Biology, 24:100–108, 2020.

[11] Matt Pennell, Oscar L Rodriguez, Corey T Watson, and Victor Greiff. The evolutionary and functional significance of germline immunoglobulin gene variation. Trends in immunology, 44(1):7–21, 2023.

[12] Oscar L Rodriguez, Xiang Qiu, Kaitlyn Shields, Christoper Dunn, Amit Singh, Mary Kaileh, Corey T Watson, and Ranjan Sen. Human genetic variation shapes the antibody repertoire across b cell development. BioRxiv, 2025.

[13] Alexandra A Fischer, Martin Corcoran, Philip JM Brouwer, Mark Chernyshev, Rebecca A Gillespie, Andrea Nicoletto, Johannes R Loeffler, James A Ferguson, Ioannis Zygouras, Pradeepa Pushparaj, et al. Genetically diverse influenza antibodies highlight the role of ig germline gene variation and inform population-comprehensive vaccine strategies. Immunity, 59(4):1123–1139, 2026.

[14] Jeong Hyun Lee, Laura Toy, Justin T Kos, Yana Safonova, William R Schief, Colin Havenar-Daughton, Corey T Watson, and Shane Crotty. Vaccine genetics of ighv1-2 vrc01-class broadly neutralizing antibody precursor naïve human b cells. NPJ vaccines, 6(1):113, 2021.

[15] Christina Yacoob, Marie Pancera, Vladimir Vigdorovich, Brian G Oliver, Jolene A Glenn, Junli Feng, D Noah Sather, Andrew T McGuire, and Leonidas Stamatatos. Differences in allelic frequency and cdrh3 region limit the engagement of hiv env immunogens by putative vrc01 neutralizing antibody precursors. Cell reports, 17(6):1560–1570, 2016.

[16] Allan C Decamp, Martin M Corcoran, William J Fulp, Jordan R Willis, Christopher A Cottrell, Daniel LV Bader, Oleksandr Kalyuzhniy, David J Leggat, Kristen W Cohen, Ollivier Hyrien, et al. Human immunoglobulin gene allelic variation impacts germline-targeting vaccine priming. npj Vaccines, 9(1):58, 2024.

[17] Yuval Avnir, Corey T Watson, Jacob Glanville, Eric C Peterson, Aimee S Tallarico, Andrew S Bennett, Kun Qin, Ying Fu, Chiung-Yu Huang, John H Beigel, et al. Ighv1-69 polymorphism modulates anti-influenza antibody repertoires, correlates with ighv utilization shifts and varies by ethnicity. Scientific reports, 6(1):20842, 2016.

[18] Pradeepa Pushparaj, Andrea Nicoletto, Daniel J Sheward, Hrishikesh Das, Xaquin Castro Dopico, Laura Perez Vidakovics, Leo Hanke, Mark Chernyshev, Sanjana Narang, Sungyong Kim, et al. Immunoglobulin germline gene polymorphisms influence the function of sars-cov-2 neutralizing antibodies. Immunity, 56(1):193–206, 2023.

[19] Martin M. Corcoran, Sanjana Narang, Mateusz Kaduk, Mark Chernyshev, Anna Färnert, Christopher Sundling, Gunilla B. Karlsson Hedestam, et al. Ultra-high-throughput igh genotyping of 25 global populations reveals population-biased allelic diversity and homozygous v and d gene deletions. Immunity, 59(4), 2026.

[20] Ayelet Peres, Amit A Upadhyay, Vered Klein, Swati Saha, Oscar L Rodriguez, Zachary M Vanwinkle, Kirti Karunakaran, Amanda Metz, William Lauer, Mark C Lin, et al. Population-level genomic analysis of immunoglobulin loci variation in rhesus macaques reveals extensive germline diversity. Immunity, 59(1):213–228, 2026.

[21] Ayelet Peres, Moriah Gidoni, Pazit Polak, and Gur Yaari. Rabhit: R antibody haplotype inference tool. Bioinformatics, 35(22):4840–4842, 2019.

[22] Zachary Sethna, Yuval Elhanati, Curtis G. Callan, Aleksandra M. Walczak, and Thierry Mora. OLGA: fast computation of generation probabilities of B- and T-cell receptor amino acid sequences and motifs. Bioinformatics, 35(17):2974–2981, 2019.

[23] Quentin Marcou, Thierry Mora, and Aleksandra M. Walczak. High-throughput immune repertoire analysis with IGoR. Nature Communications, 9:561, 2018.

[24] Branden J Olson and Frederick A Matsen IV. The bayesian optimist’s guide to adaptive immune receptor repertoire analysis. Immunological reviews, 284(1):148–166, 2018.

[25] Thomas Konstantinovsky, Ayelet Peres, Pazit Polak, and Gur Yaari. An unbiased comparison of immunoglobulin sequence aligners. Briefings in bioinformatics, 25(6):bbae556, 2024.

[26] Ann J Feeney, Alan Tang, and Kisani M Ogwaro. B-cell repertoire formation: role of the recombination signal sequence in non-random v segment utilization. Immunological reviews, 175(1):59–69, 2000.

[27] Amy L Kenter, Corey T Watson, and Jan-Hendrik Spille. Igh locus polymorphism may dictate topological chromatin conformation and v gene usage in the ig repertoire. Frontiers in Immunology, 12:682589, 2021.

[28] Amy L Kenter, Saurabh Priyadarshi, and Ellen B Drake. Locus architecture and rag scanning determine antibody diversity. Trends in immunology, 44(2):119–128, 2023.

[29] Khalid H Bhat, Saurabh Priyadarshi, Sarah Naiyer, Xinyan Qu, Hammad Farooq, Eden Kleiman, Jeffery Xu, Xue Lei, Jose F Cantillo, Robert Wuerffel, et al. An igh distal enhancer modulates antigen receptor diversity by determining locus conformation. Nature communications, 14(1):1225, 2023.

[30] Wen-Wei Liao, Mobin Asri, Jana Ebler, Daniel Doerr, Marina Haukness, Glenn Hickey, Shuangjia Lu, Julian K. Lucas, Jean Monlong, Haley J. Abel, et al. A draft human pangenome reference. Nature, 617(7960):312–324, 2023.

[31] The 1000 Genomes Project Consortium. A global reference for human genetic variation. Nature, 526(7571):68–74, 2015.

[32] Aviv Omer, Or Shemesh, Ayelet Peres, Pazit Polak, Adrian J Shepherd, Corey T Watson, Scott D Boyd, Andrew M Collins, William Lees, and Gur Yaari. Vdjbase: an adaptive immune receptor genotype and haplotype database. Nucleic acids research, 48(D1):D1051–D1056, 2020.

[33] William D. Lees, Ayelet Peres, Vered Klein, Naama Amos, Uddalok Jana, Eric Engelbrecht, Zachary Vanwinkle, Yaniv Malach, Thomas Konstantinovsky, Pazit Polak, Corey T. Watson, and Gur Yaari. The current landscape of adaptive immune receptor genomic and repertoire data: OGRDB and VDJbase. Nucleic Acids Research, 54(D1):D932–D937, 2026.

[34] Lindsay G Cowell, Scott Christley, Felix Breden, Kevin A Burns, Brian D Corrie, William D Lees, James A Overton, Bjoern Peters, Eve Richardson, Krishna M Roskin, et al. The adaptive immune receptor repertoire knowledge commons: an invitation to the community. Cell Systems, 16(9), 2025.

[35] 1000 Genomes Project Consortium et al. A map of human genome variation from population scale sequencing. Nature, 467(7319):1061, 2010.

[36] Haoyu Cheng, Gregory T Concepcion, Xiaowen Feng, Haowen Zhang, and Heng Li. Haplotype-resolved de novo assembly using phased assembly graphs with hifiasm. Nature methods, 18(2):170–175, 2021.

[37] Uddalok Jana, Zach Vanwinkle, and Corey Watson. WASP (Workflow for Antibody Sequence Profiling), March 2026. Workflow.

[38] Ayelet Peres, William D Lees, Oscar L Rodriguez, Noah Y Lee, Pazit Polak, Ronen Hope, Meirav Kedmi, Andrew M Collins, Mats Ohlin, Steven H Kleinstein, Corey T Watson, and Gur Yaari. IGHV allele similarity clustering improves genotype inference from adaptive immune receptor repertoire sequencing data. Nucleic Acids Research, page gkad603. 08 2023.

[39] Eric Engelbrecht, Oscar L Rodriguez, and Corey T Watson. Addressing technical pitfalls in pursuit of molecular factors that mediate immunoglobulin gene regulation. The Journal of Immunology, 213(5):651–662, 2024.

[40] Cathrine Scheepers, Ram K Shrestha, Bronwen E Lambson, Katherine JL Jackson, Imogen A Wright, Dshanta Naicker, Mark Goosen, Leigh Berrie, Arshad Ismail, Nigel Garrett, et al. Ability to develop broadly neutralizing hiv-1 antibodies is not restricted by the germline ig gene repertoire. The Journal of Immunology, 194(9):4371–4378, 2015.

[41] Corey T Watson, Karyn Meltz Steinberg, Tina A Graves, Rene L Warren, Maika Malig, Jacqueline Schein, Richard K Wilson, Robert A Holt, Evan E Eichler, and Felix Breden. Sequencing of the human ig light chain loci from a hydatidiform mole bac library reveals locus-specific signatures of genetic diversity. Genes & Immunity, 16(1):24–34, 2015.

[42] David G Schatz, Marjorie A Oettinger, and Mark S Schlissel. V (d) j recombination: molecular biology and regulation. Annual review of immunology, 10(1):359–383, 1992.

[43] Michael J Difilippantonio, Catherine J McMahan, Quinn M Eastman, Eugenia Spanopoulou, and David G Schatz. Rag1 mediates signal sequence recognition and recruitment of rag2 in v (d) j recombination. Cell, 87(2):253–262, 1996.

[44] Dik C Van Gent, Dale A Ramsden, and Martin Gellert. The rag1 and rag2 proteins establish the 12/23 rule in v (d) j recombination. Cell, 85(1):107–113, 1996.

[45] Alfred Ian Lee, Sebastian D Fugmann, Lindsay G Cowell, Leon M Ptaszek, Garnett Kelsoe, and David G Schatz. A functional analysis of the spacer of v (d) j recombination signal sequences. PLoS biology, 1(1):e1, 2003.

[46] Fumihiko Matsuda, Kazuo Ishii, Patrice Bourvagnet, Kei-ichi Kuma, Hidenori Hayashida, Takashi Miyata, and Tasuku Honjo. The complete nucleotide sequence of the human immunoglobulin heavy chain variable region locus. The Journal of experimental medicine, 188(11):2151–2162, 1998.

[47] Oscar L Rodriguez, William S Gibson, Tom Parks, Matthew Emery, James Powell, Maya Strahl, Gintaras Deikus, Kathryn Auckland, Evan E Eichler, Wayne A Marasco, et al. A novel framework for characterizing genomic haplotype diversity in the human immunoglobulin heavy chain locus. Frontiers in immunology, 11:2136, 2020.

[48] Marie J Kidd, Katherine JL Jackson, Scott D Boyd, and Andrew M Collins. Dj pairing during vdj recombination shows positional biases that vary among individuals with differing ighd locus immunogenotypes. The Journal of Immunology, 196(3):1158–1164, 2016.

[49] Ryan Haryadi, Steven Ho, Yee Jiun Kok, Helen X Pu, Lu Zheng, Natasha A Pereira, Bin Li, Xuezhi Bi, Lin-Tang Goh, Yuansheng Yang, et al. Optimization of heavy chain and light chain signal peptides for high level expression of therapeutic antibodies in cho cells. PloS one, 10(2):e0116878, 2015.

[50] Hajar Owji, Navid Nezafat, Manica Negahdaripour, Ali Hajiebrahimi, and Younes Ghasemi. A comprehensive review of signal peptides: Structure, roles, and applications. European journal of cell biology, 97(6):422–441, 2018.

[51] Oscar L Rodriguez, Andrew J Sharp, and Corey T Watson. Limitations of lymphoblastoid cell lines for establishing genetic reference datasets in the immunoglobulin loci. Plos one, 16(12):e0261374, 2021.

[52] Mao-Jan Lin, Ben Langmead, and Yana Safonova. Igloo enables comprehensive analysis and assembly of immunoglobulin heavy-chain loci in lymphoblastoid cell lines using pacbio high-fidelity reads. Cell Reports Methods, 5(5), 2025.

[53] Achmad Dimas Cahyaning Furqon, Leah W Roberts, and Michael B Hall. Efficient downsampling of genome alignments with rasusa. Gigabyte, 2026:gigabyte180–0, 2026.

[54] Michael B. Hall. Rasusa: Randomly subsample sequencing reads to a specified coverage. Journal of Open Source Software, 7(69):3941, 2022.

