## Supplementary Information for "Population-scale immunoglobulin genomics links inherited variation to antibody repertoire organization"

Ayelet Peres[1,*], Uddalok Jana[2,*], Oscar L. Rodriguez[2], Zachary M. Vanwinkle[2], Eric Engelbrecht[2], William S Gibson[2], Kaitlyn Shields[2], Brandon Croslin[2], Steven Schultze[2], Chandrima Bharadwaj[2], Connor Murray[2], William D. Lees[3], Melissa L. Smith[2,†,§], Corey T. Watson[2,†,§] and Gur Yaari[1,†,§]

[1]Department of Pathology, Yale School of Medicine, New Haven, CT, USA
[2]Department of Biochemistry and Molecular Genetics, University of Louisville, Louisville, KY, USA
[3]Clareo Biosciences, Louisville, KY 40202, United States
[*]These authors contributed equally to this work
[†]These authors jointly supervised this work

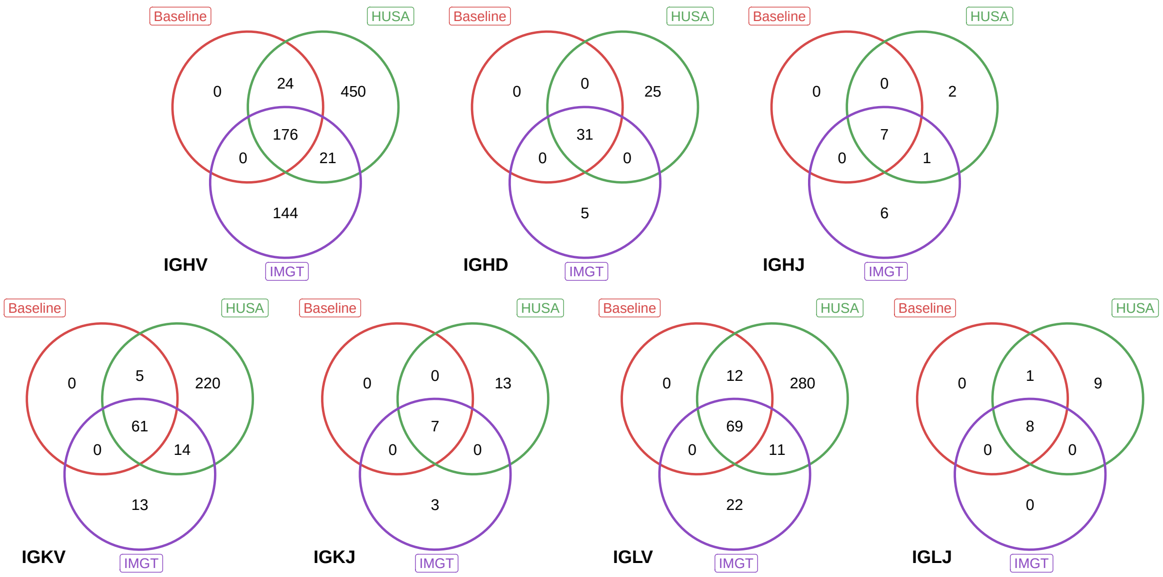


Figure 1: **HUSA expansion relative to baseline and IMGT allele sets.** Venn diagrams show overlap among coding-region allele sequences represented in the baseline reference set, HUSA, and IMGT for IGHV, IGHD, IGHJ, IGKV, IGKJ, IGLV, and IGLJ. Overlap was computed by exact coding-region sequence identity within each gene segment, with the IMGT set restricted to functional and ORF alleles of genes represented in HUSA. Numbers indicate the count of allele sequences in each intersection. The baseline set was constructed from the OGRDB human immunoglobulin reference supplemented with additional IMGT alleles. This comparison summarizes how HUSA relates to the annotation seed reference and to IMGT across immunoglobulin gene segments.

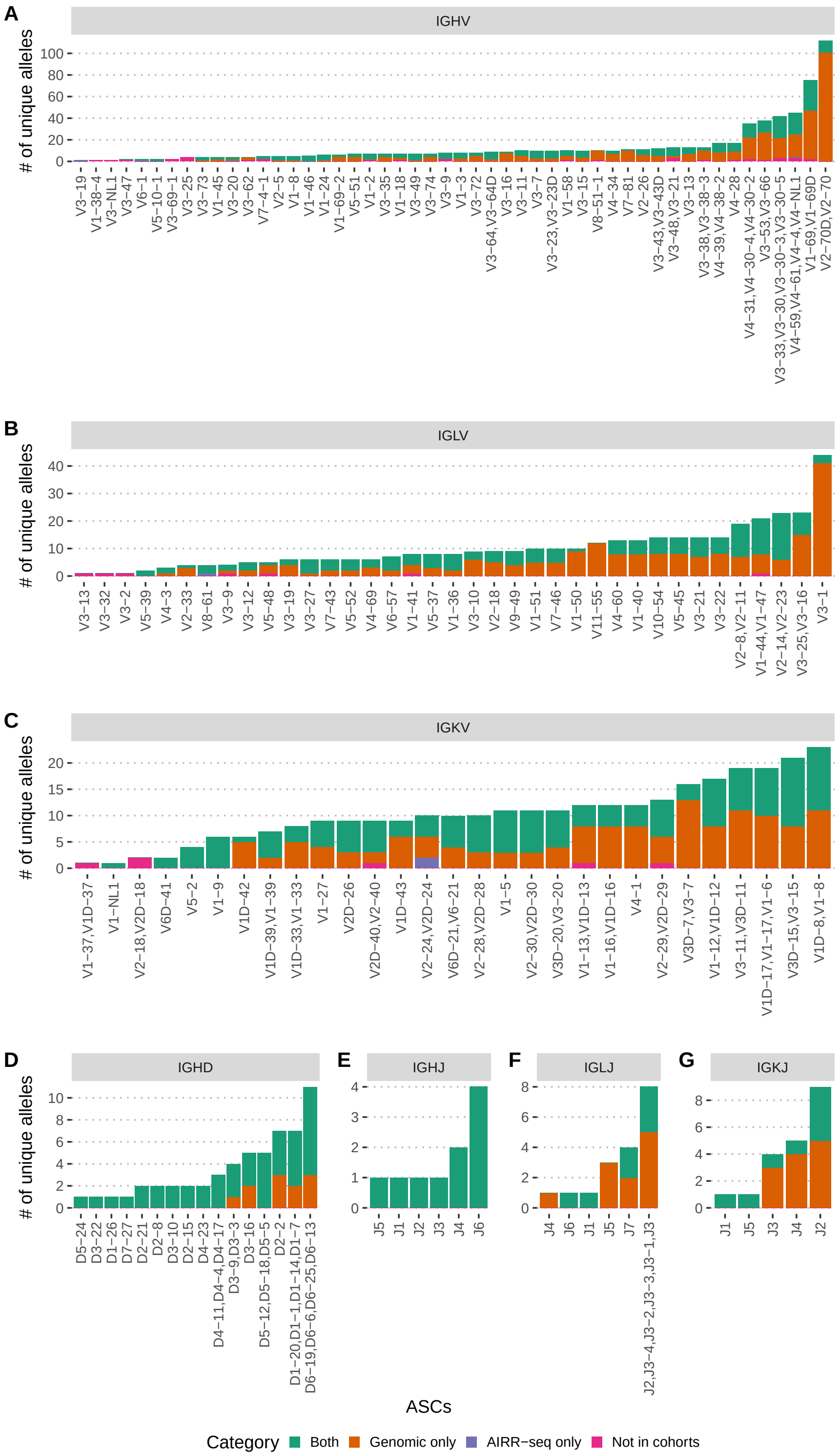


Figure 2: **Distribution of unique alleles across allele similarity clusters.** (A–G) Stacked bar plots showing the number of unique alleles assigned to each allele similarity cluster (ASC) for IGHV (A), IGLV (B), IGKV (C), IGHD (D), IGHJ (E), IGLJ (F), and IGKJ (G). ASCs are labeled by the genes represented in each cluster, with overlapping ASCs merged and displayed using combined gene labels. Color indicates whether alleles were observed in both genomic and AIRR-seq cohorts, genomic data only, AIRR-seq data only, or were not observed in the analyzed cohorts.

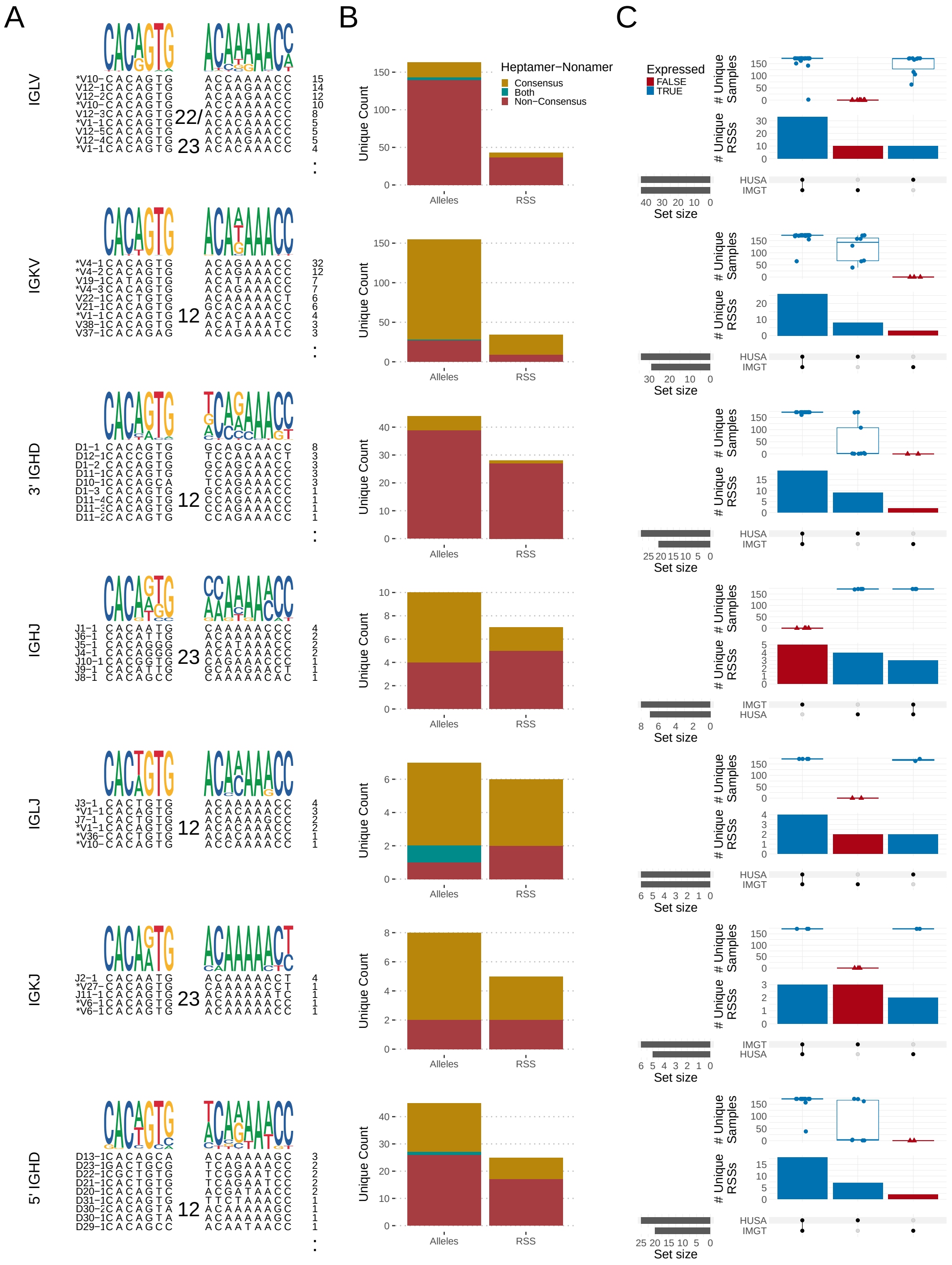


Figure 3: **Diversity and overlap of recombination signal sequences (RSS) across immunoglobulin loci.** (A) Sequence logos of the heptamer and nonamer motifs for each gene segment, weighted by the number of unique allele-RSS sequence pairs in expressed alleles of DS3. The number of unique allele-RSS pairs per motif is shown to the right of each sequence. Asterisks denote heptamer-nonamer combinations also observed in other loci. (B) Bar plots showing the number of unique germline alleles and associated RSS variants per locus. (C) UpSet plots illustrating intersections between RSS variants identified in HUSA and those present in IMGT. Horizontal bars show the number of distinct RSSs in each source, and vertical bars represent shared or source-specific RSS variants. Boxplots show the number of individuals carrying each RSS variant. Color indicates whether the RSS variant was linked to an allele observed in expressed repertoire data.

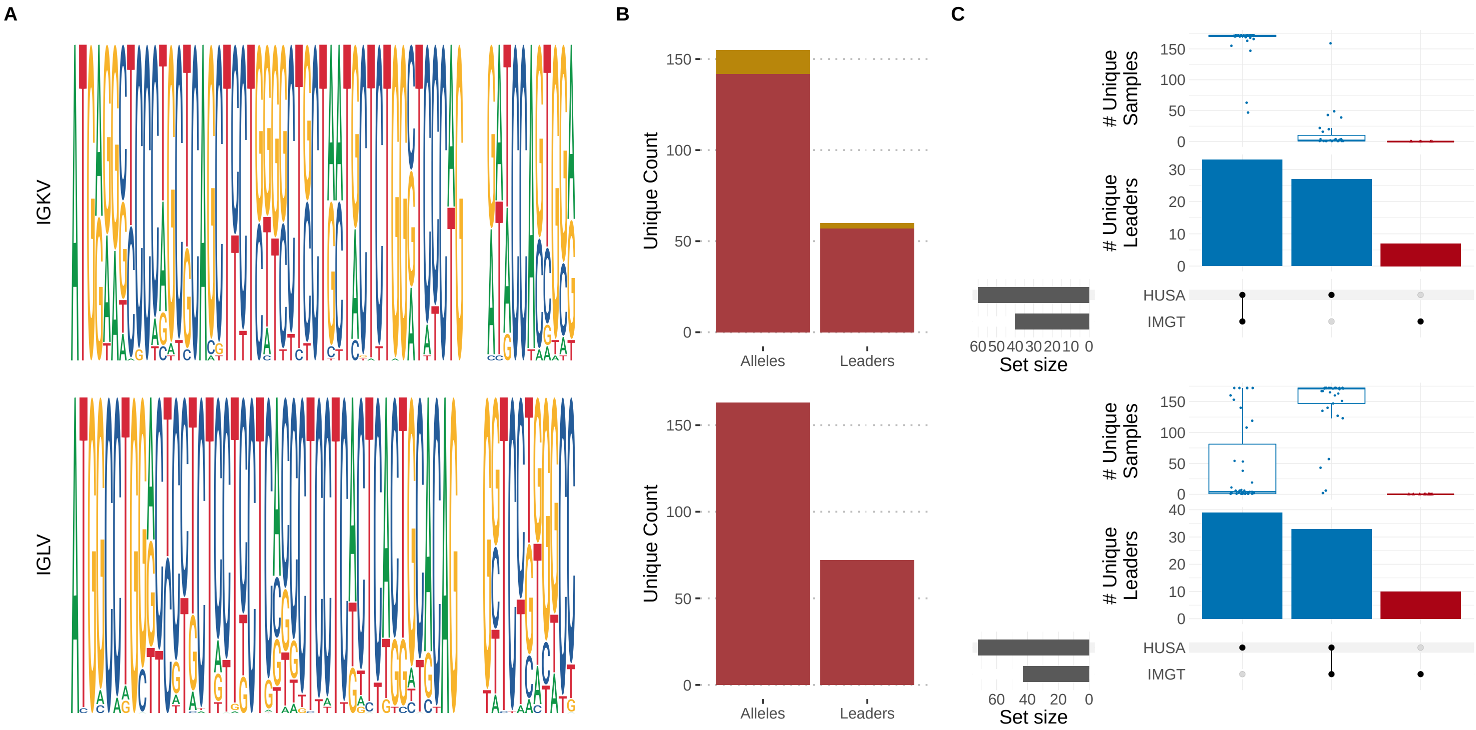


Figure 4: **Leader sequence diversity and HUSA-IMGT leader overlap in IGKV and IGLV.** (A) Sequence logos of leader sequences for IGKV and IGLV genes, based on leader sequences from expressed alleles of DS3. (B) Counts of unique alleles and leader sequences for each V-gene locus, stratified by leader consensus status. (C) UpSet plots comparing leader sequences represented in HUSA and IMGT. Bars show the number of unique leader sequences in each source-specific or shared intersection, boxplots show the number of samples carrying each leader sequence, and color indicates whether the associated allele was observed in expressed repertoire data.

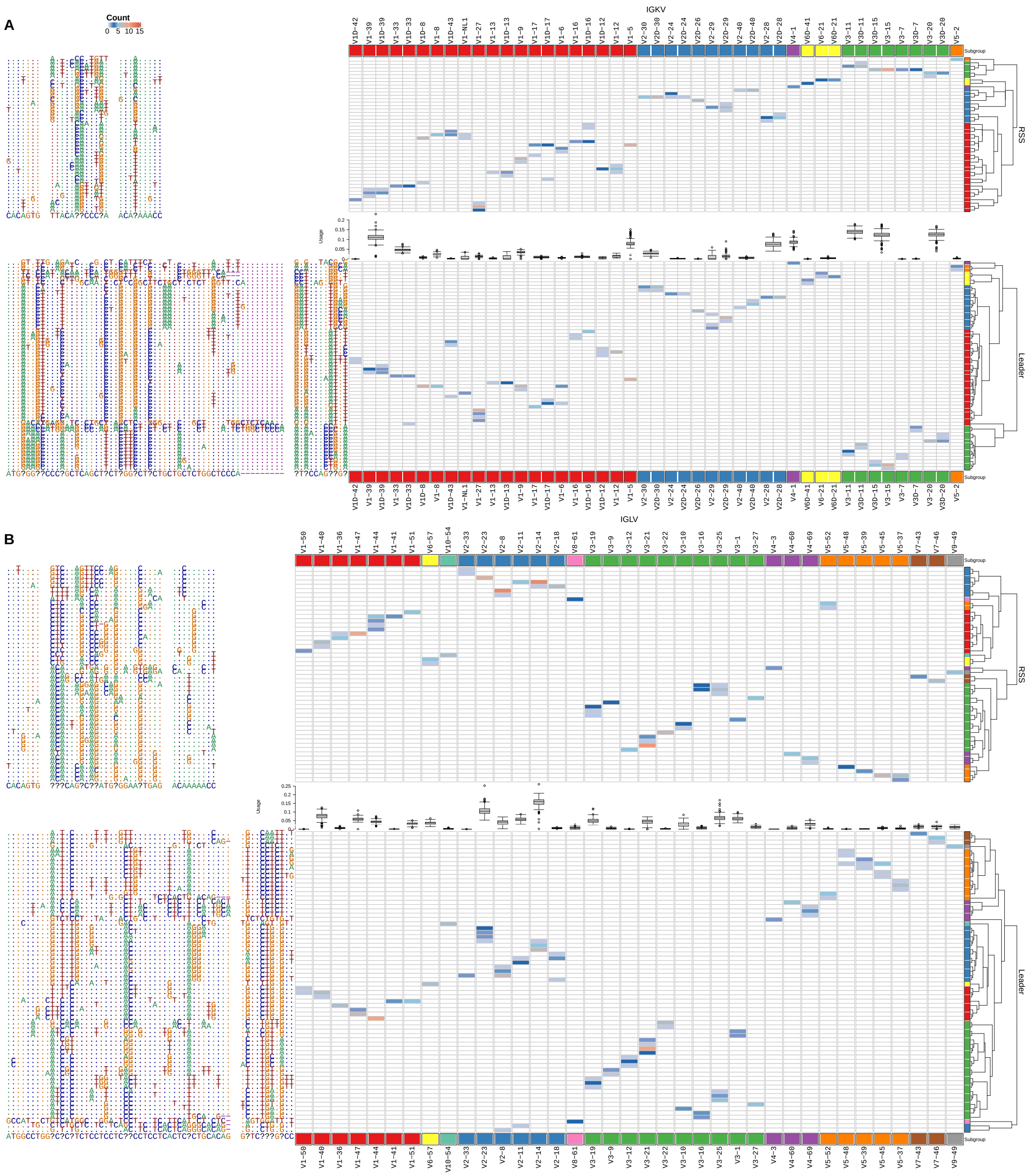

Figure 5: **Coding-region, RSS, leader, and repertoire-usage variation in IGKV and IGLV ASCs.** (A) Heatmaps for IGKV ASCs showing the relationship between coding-region ASCs and their corresponding RSS sequences, relative usage in expressed repertoires, and full-length leader sequence variants. Cell color denotes the number of alleles with each variant combination. Rows in the RSS and leader panels are ordered by hierarchical clustering, and columns are ordered by chromosomal position within subgroup. (B) Same analysis for IGLV ASCs.

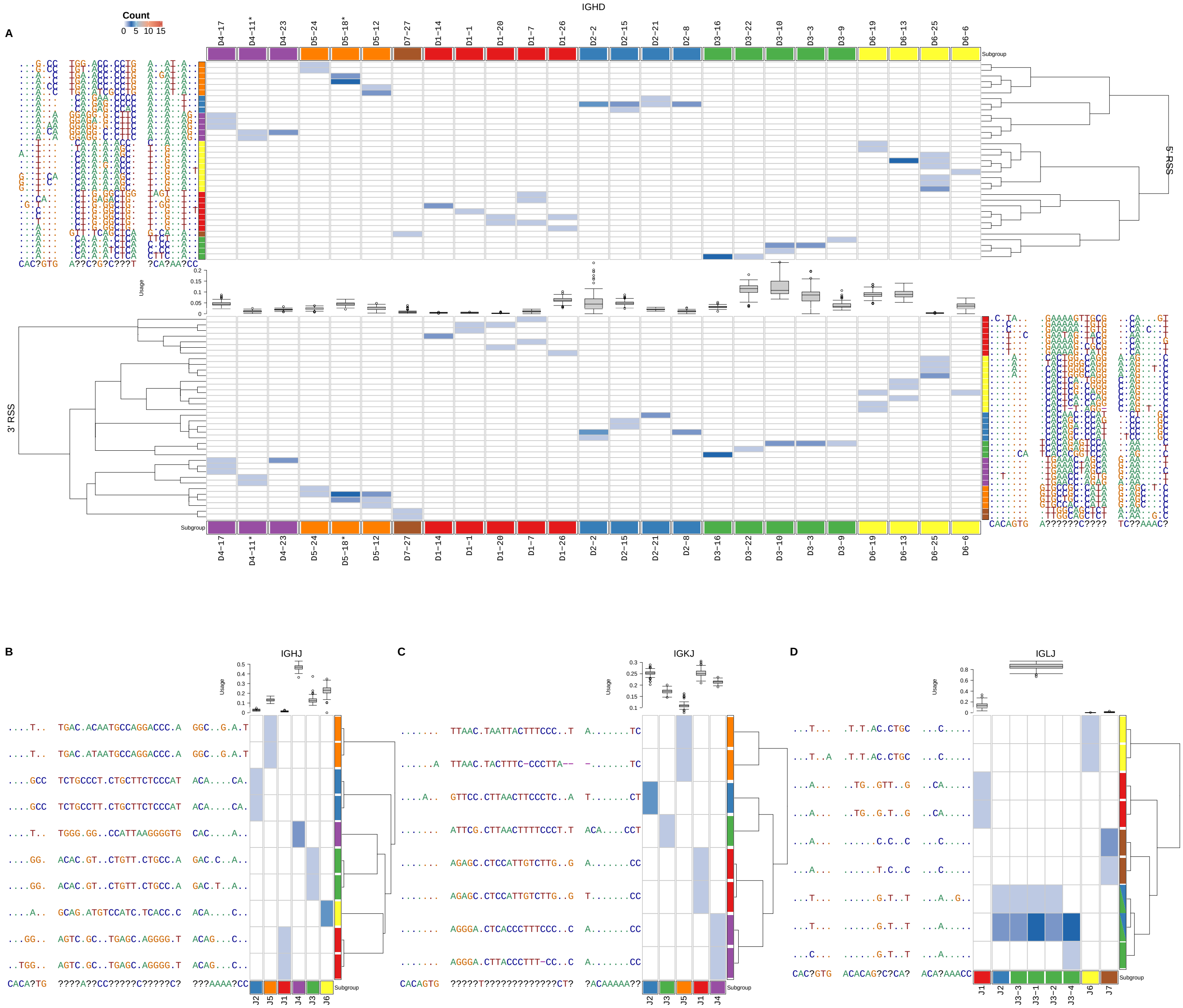


Figure 6: **RSS variation and relative usage across D and J gene segments.** (A) IGHD ASCs showing the relationship between coding-region variation, 5′ RSS sequences, relative usage in expressed repertoires, and 3′ RSS sequences. Cell color denotes the number of alleles with each variant combination. Rows are ordered by hierarchical clustering, and columns are ordered by chromosomal position. (B) IGHJ ASCs showing relative usage and RSS sequence variation. (C) IGKJ ASCs analyzed as in (B). (D) IGLJ ASCs analyzed as in (B).

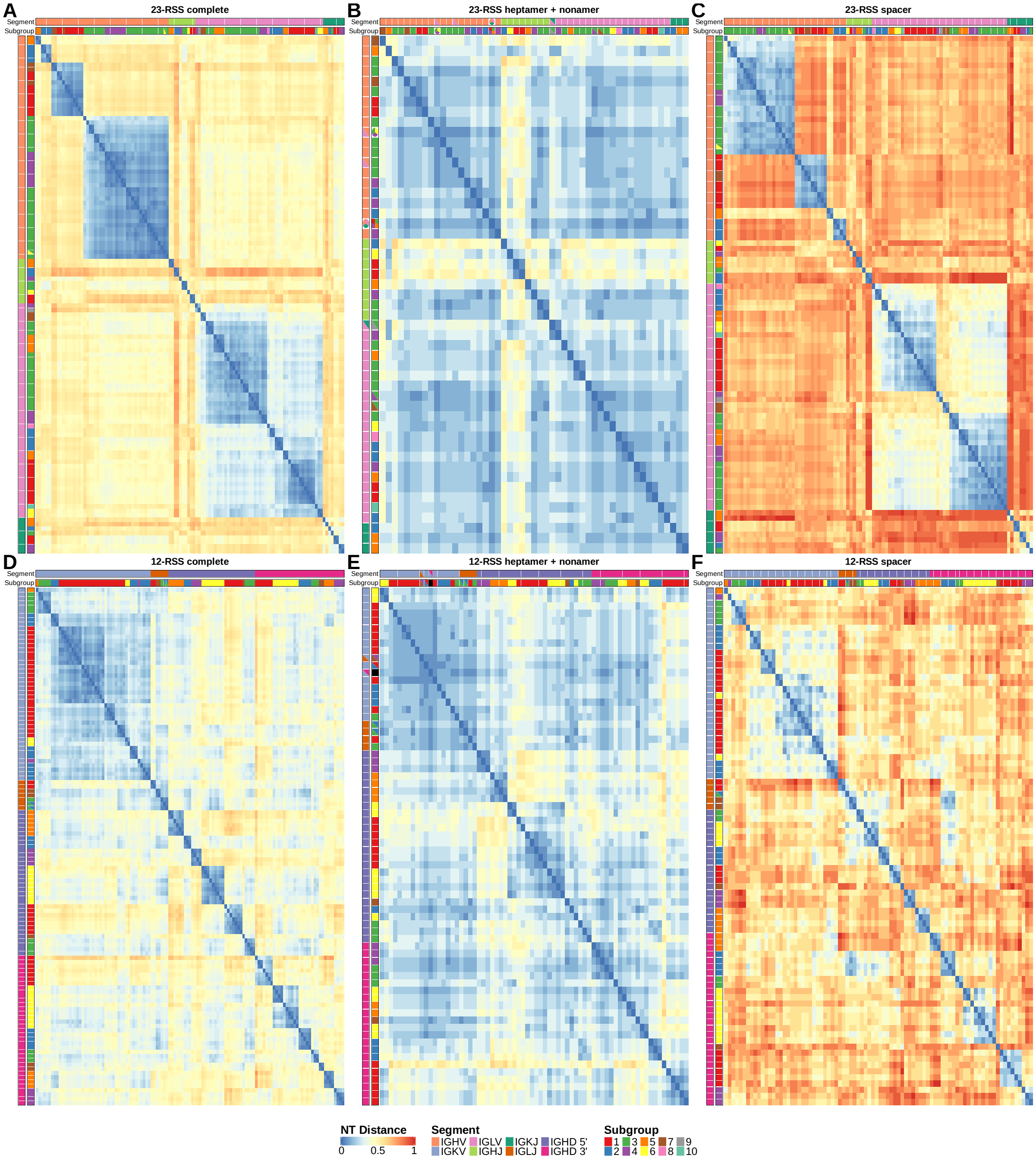


Figure 7: **RSS conservation and coding-noncoding sequence-distance concordance support partial co-organization of IG coding and linked regulatory features.** (A-C) Nucleotide-distance heatmaps for RSSs with 23-bp spacers: complete RSS (A), heptamer plus nonamer (B), and spacer-only distances (C). Segment and subgroup annotations are shown on rows and columns, with hierarchical clustering applied within segment groups. (D-F) Nucleotide-distance heatmaps for RSSs with 12-bp spacers: complete RSS (D), heptamer plus nonamer (E), and spacer-only distances (F), displayed as in (A-C).

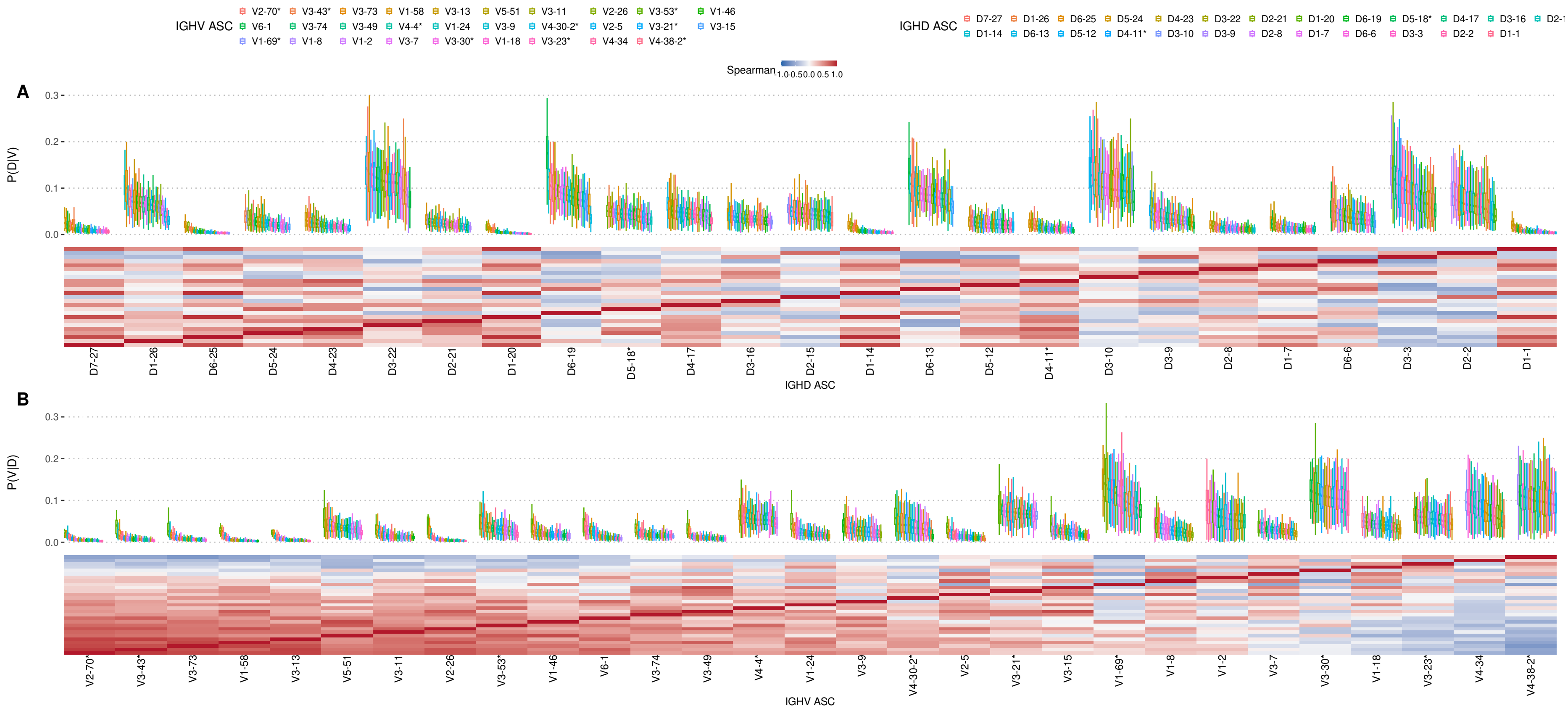


Figure 8: **Conditional IGH V–D usage relationships across allele similarity clusters.** Alleles were grouped into allele similarity clusters (ASCs) to reduce ambiguity from highly similar coding-region sequences. Conditional usage probabilities were computed at the ASC level, and heatmaps show Spearman correlations between the resulting segment-usage rank orders. Axis labels ending in an asterisk denote shortened ASC labels; full gene membership for these labels is provided in Sup. Table 1. (A) $P(D \mid V)$, showing D-segment usage conditional on IGHV ASC. Boxplots show individual-level conditional usage distributions, and the heatmap shows Spearman correlations between IGHD ASC usage profiles. (B) $P(V \mid D)$, showing V-segment usage conditional on IGHD ASC. Boxplots show individual-level conditional usage distributions, and the heatmap shows Spearman correlations between IGHV ASC usage profiles.

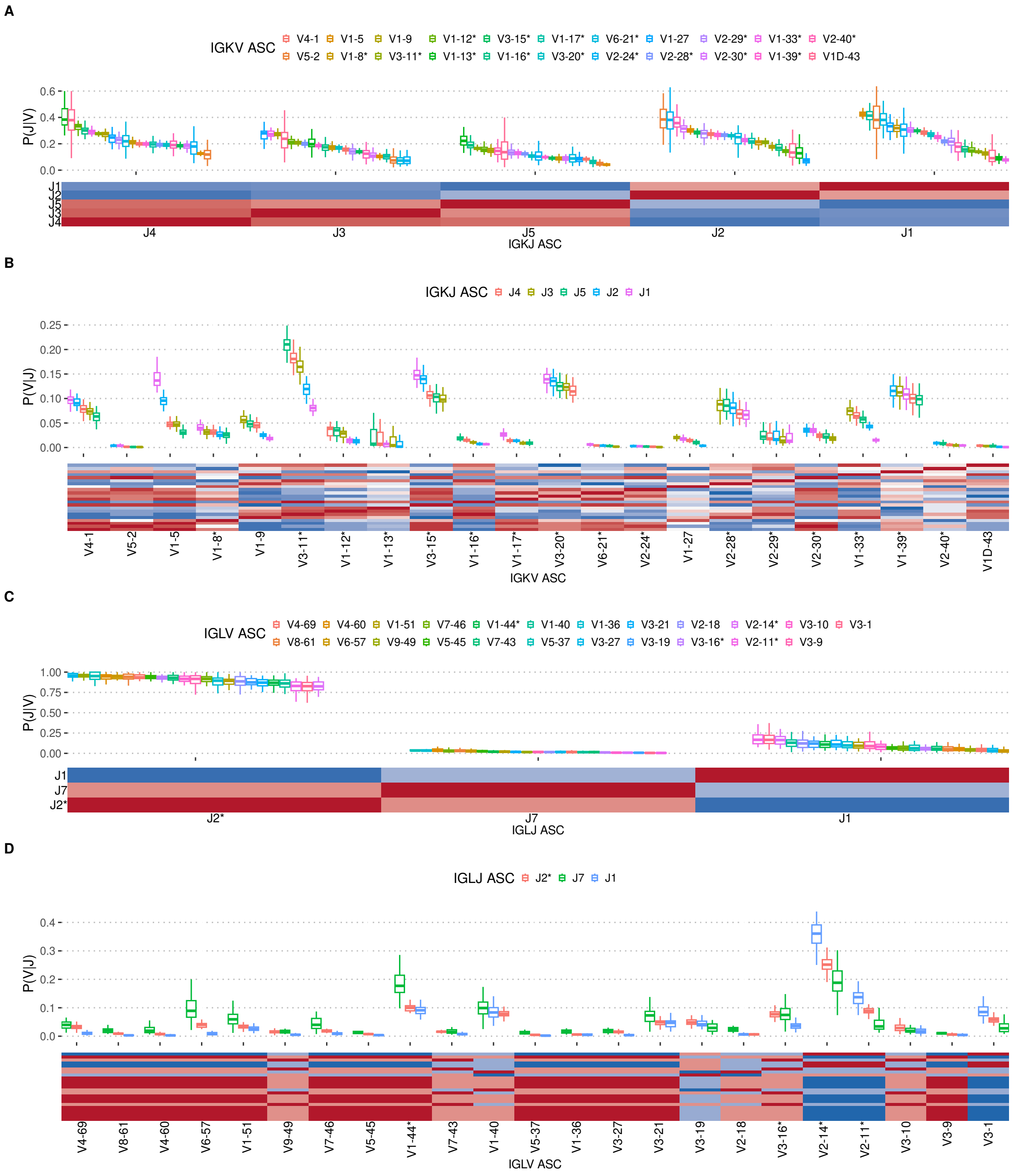


Figure 9: **Conditional IGK and IGL V–J usage relationships across allele similarity clusters.** Conditional usage probabilities were computed at the ASC level, and heatmaps show Spearman correlations between segment-usage profiles. Asterisks denote shortened ASC labels; full membership is provided in Sup. Table 1. (A-B) IGK $P(J \mid V)$ and $P(V \mid J)$, respectively. (C-D) IGL $P(J \mid V)$ and $P(V \mid J)$, respectively. Boxplots show individual-level conditional usage distributions.

Table 1: **Shortened ASC labels used in conditional segment-usage figures.** Plot-axis labels ending in an asterisk denote allele similarity clusters (ASCs) for which multiple genes were collapsed into a shortened label for visualization in Figure 4 and Sup. Figure 8. The full set of genes represented by each shortened ASC label is listed.

| **Axis** | **Plot axis label** | **ASC genes** |
|---|---|---|
| IGHD | D4-11* | D4-11/4-4 |
| IGHD | D5-18* | D5-18/5-5 |
| IGHV | V1-69* | V1-69/1-69D |
| IGHV | V2-70* | V2-70/2-70D |
| IGHV | V3-21* | V3-21/3-48 |
| IGHV | V3-23* | V3-23/3-23D |
| IGHV | V3-30* | V3-30/3-30-3/3-30-5/3-33 |
| IGHV | V3-43* | V3-43/3-43D |
| IGHV | V3-53* | V3-53/3-66 |
| IGHV | V4-30-2* | V4-30-2/4-30-4/4-31 |
| IGHV | V4-38-2* | V4-38-2/4-39 |
| IGHV | V4-4* | V4-4/4-59/4-61/4-NL1 |
| IGKV | V1-8* | V1-8/1D-8 |
| IGKV | V1-12* | V1-12/1D-12 |
| IGKV | V1-13* | V1-13/1D-13 |
| IGKV | V1-16* | V1-16/1D-16 |
| IGKV | V1-17* | V1-17/1-6/1D-17 |
| IGKV | V1-33* | V1-33/1D-33 |
| IGKV | V1-39* | V1-39/1D-39 |
| IGKV | V2-24* | V2-24/2D-24 |
| IGKV | V2-28* | V2-28/2D-28 |
| IGKV | V2-29* | V2-29/2D-29 |
| IGKV | V2-30* | V2-30/2D-30 |
| IGKV | V2-40* | V2-40/2D-40 |
| IGKV | V3-11* | V3-11/3D-11 |
| IGKV | V3-15* | V3-15/3D-15 |
| IGKV | V3-20* | V3-20/3D-20 |
| IGKV | V6-21* | V6-21/6D-21 |
| IGLV | V1-44* | V1-44/1-47 |
| IGLV | V2-11* | V2-11/2-8 |
| IGLV | V2-14* | V2-14/2-23 |
| IGLV | V3-16* | V3-16/3-25 |
| IGLJ | J2* | J2/3-1/3-2/3-3/3-4 |

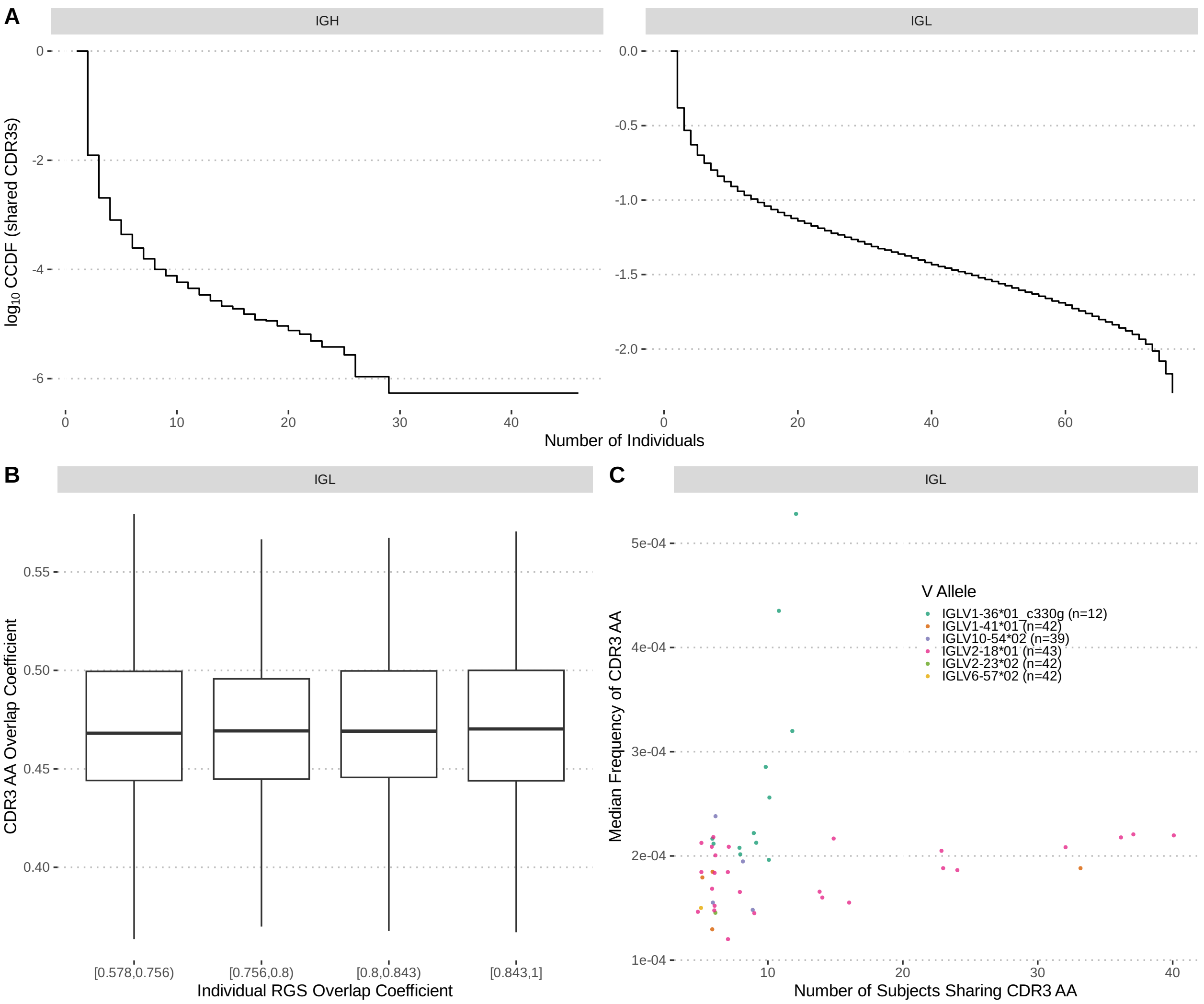


Figure 10: **Additional CDR3 amino acid sharing analyses across expressed repertoires in DS3.** (A) Complementary cumulative distribution functions (CCDFs) of exact CDR3 amino acid (AA) sequences shared across individuals, shown by locus. The x-axis shows the number of individuals sharing a CDR3 AA sequence, and the y-axis shows the log10-transformed cumulative probability of sequences shared by at least that number of individuals. (B) Relationship between individual RGS overlap and CDR3 AA repertoire overlap across pairs of individuals, shown by locus. RGS overlap was calculated from PIgLET-inferred repertoire genotype sets, and repertoire overlap was calculated from exact CDR3 AA sets. (C) V allele-specific shared CDR3 AA sequences for additional light-chain analyses. Each point represents an exact CDR3 AA sequence observed in at least five individuals and assigned to a single V allele. The x-axis shows the number of subjects sharing the CDR3 AA sequence, and the y-axis shows its median within-individual frequency. Only V alleles observed in no more than 60% of individuals in the corresponding locus were included.
